# A Machine Learning Approach Towards the Differentiation Between Interoceptive and Exteroceptive Attention

**DOI:** 10.1101/2022.06.10.495649

**Authors:** Zoey X. Zuo, Cynthia J. Price, Norman A. S. Farb

**Author notes:** Corresponding Author: Zoey X. Zuo, Graduate Department of Psychological Clinical Science, University of Toronto Scarborough, 1265 Military Trail, Scarborough, ON M1C 1A4, Canada.

## Abstract

Interoception, the representation of the body’s internal state, plays a central role in emotion, motivation, and wellbeing. Interoceptive attention is qualitatively different from attention to the external senses and may recruit a distinct neural system, but the neural separability of interoceptive and exteroceptive attention is unclear. We used a machine learning approach to classify neural correlates of interoceptive and exteroceptive attention in a randomized control trial of interoceptive training (MABT). Participants in the training and control groups attended fMRI assessment before and after an 8-week intervention period (N = 44 scans). The imaging paradigm manipulated attention targets (breath vs. visual stimulus) and reporting demands (active reporting vs. passive monitoring). Machine learning models achieved high accuracy in distinguishing between interoceptive and exteroceptive attention using both in-sample and more stringent out-of-sample tests. We then explored the potential of these classifiers in “reading out” mental states in a sustained interoceptive attention task. Participants were classified as maintaining an active reporting state for only ∼90s of each 3-minute sustained attention period. Within this active period, interoceptive training enhanced participants’ ability to sustain interoceptive attention. These findings demonstrate that interoceptive and exteroceptive attention engage reliable and distinct neural networks; machine learning classifiers trained on this distinction show promise for assessing the stability of interoceptive attention, with implications for the future assessment of mental health and treatment response.

## Introduction

Interoception, the sense of the body’s internal state, is widely regarded as a foundation for emotion (Wiens, 2005), motivation (Craig, 2003), intuition (Dunn et al., 2010), and wellbeing (Tsakiris & Critchley, 2016). While interoceptive signals such as respiration, heart rate, temperature or hunger are processed automatically to promote homeostasis in the body (Craig, 2002; Gu & Fitzgerald, 2014), these signals also serve as a foundation for feeling states that guide consciously coordinated behavior (Strigo & Craig, 2016). As such, interoceptive function has recently become the target of psychological theory casting the interoceptive sense at the heart of health and disease, characterizing interoception as affectively privileged and distinct from the exteroceptive senses of vision, hearing, taste, smell, and touch (Farb et al., 2015; Khalsa et al., 2018; Quadt et al., 2018).

Considerable research supports the characterization of interoception as distinct from the exteroceptive senses. Interoception is supported by a dedicated neuroanatomical pathway, with signals transmitted via sense-receptor C-fiber afferents along the spinal cord to the brainstem and thalamus before reaching the cerebral cortex (Craig, 2002; Critchley & Harrison, 2013). Functionally, interoceptive attention appears to engage the ventromedial thalamus and right posterior insula (Farb et al., 2013), which serves as the primary interoceptive cortex (Craig, 2002; Flynn, 1999). From a phenomenological perspective, interoception is also seen as a distinct type of experience or ‘sixth sense’ that provides a sense of presence (Farb et al., 2015) or affective tone (Barrett & Quigley, 2021), affording living organisms an affectively embodied perspective (Smith, 2022).

Yet while interoception seems qualitatively distinct from exteroception, it is unknown whether is supported by a distinct attentional network. A set of frontoparietal brain regions constitute a well-validated Dorsal Attention Network (DAN), which is broadly involved in the regulation of perceptual attention (Dixon et al., 2018; Szczepanski et al., 2013). It is plausible that the DAN also supports attention for interoceptive signals, given that interoceptive experiences are usually multimodal combinations of interoceptive and exteroceptive afferents (Khalsa et al., 2009; Quigley et al., 2021). Furthermore, despite evidence that the anterior insula facilitates subjective access to body sensations, this integration may not be modality specific (Craig, 2009; Critchley et al., 2004; Gu et al., 2013). Instead, the anterior insula seems to integrate both interoceptive and exteroceptive signals (Medford & Critchley, 2010; Seth et al., 2012) and serves as the sensory/afferent hub of the salience network (Seeley, 2019), which could then provide a common ‘neural code’ to higher cognitive processes such as the regulation of perceptual attention supported by the DAN.

While primary sensory cortices clearly engage distinct neural populations, subsequent evaluation and regulation of these signals following integration are in polymodal association cortices, as suggested by recent work demonstrating that both olfaction and vision share a common affective code in the ventromedial prefrontal cortex (Chikazoe et al., 2014). These findings are also supported by prominent theories of mindfulness (Brown & Cordon, 2009) and psychological appraisal (Ellsworth, 2013; Siemer & Reisenzein, 2007), which both place affective responses as subsequent to sense perception of its internal/external origin. The regulation of interoceptive signals has important clinical implications for conditions such anxiety (Domschke et al., 2010), obesity (Herbert & Pollatos, 2014), chronic pain (Pollatos et al., 2012), and body perception (Tsakiris et al., 2011). As such, determining whether neural representations of interoceptive attention are distinct from exteroceptive attention remains an important and largely unaddressed area of research.

Machine learning approaches offer a novel approach to resolving the question of interoceptive attention’s neural distinctiveness. Time-series data from functional neuroimaging, such as fMRI, is often analyzed using machine learning algorithms to identify meaningful brain patterns (e.g., Haxby, 2012; Norman et al., 2006) such as predicting which traumatic film scenes became intrusive memories (Clark et al., 2014). Compared to univariate methods, machine learning maximizes the potential to test for meaningful distinctions in fMRI data (Davatzikos, 2019). If applied to the classification of interoceptive attention, a machine learning approach could shed light on the separability of interoception’s neural correlates from the attention to the external senses. Supporting the feasibility of such classification, Weng et al., (2020) documented the ability for machine learning to distinguish between different internal targets of attention (i.e., focus on the breath, the self, or mind-wandering) using a logistic regression model, a classification scheme which was promising for understanding the effects of meditation expertise on attentional stability.

The successful classification between interoceptive and exteroceptive attention could in turn inform clinical science. Compromised interoceptive functioning has long been associated with general health and mental health concerns (see Khalsa et al., 2018, for a review). For example, major depressive disorder (MDD) symptoms have been associated with dysregulated interoceptive processes (Dunn et al., 2007; Terhaar et al., 2012), and individuals with depression exhibit lower sensitivity to interoceptive signals (Furman et al., 2013). Symptoms that underlie panic disorder, somatic symptom disorders, posttraumatic stresdisorder (PTSD), and substance use disorders, have all been associated with interoceptive dysfunction(Barsky et al., 2001; Glenn et al., 2016; Naqvi & Bechara, 2010). Many mindfulness-based clinical interventions have demonstrated that increases in interoceptive awareness is concomitant with reduced symptoms of distress, suggesting that interoception is amenable to training and that its improvement brings about health benefits (e.g., Mindfulness-Based Cognitive Therapy for chronic pain and concurrent depression (de Jong et al., 2016), Dialectical Behavior Therapy for generalized anxiety disorder (Navarro-Haro et al., 2019), and Mindful Awareness in Body-oriented Therapy (MABT) for substance use disorder treatment (Price et al., 2012, 2019, 2020). Of these approaches, MABT (Price & Hooven, 2018) appears to improve interoceptive attention over time and with practice, such that participants gain greater capacity to access, and sustain their attention to interoceptive experience. If machine learning models can successfully distinguish between neural representations of interoception and exteroception, they could be applied to assess the impact of interoceptive training, clarifying the relationship between changes in interoceptive function and wellbeing.

Accordingly, we applied a machine learning approach to a novel fMRI paradigm that scaffolded interoceptive and exteroceptive attention. The study was conducted in the context of a well-validated mindfulness-based intervention (MABT), which was particularly appropriate for this study due to its unique focus on teaching fundamental skills critical to identifying, accessing, sustaining, and appraising signals that arise within the body (Price & Hooven, 2018). We then applied the classification model to explore the effects of interoceptive training on sustained interoceptive attention. Specifically, we had four aims in this study:

1. Distinguish between the neural patterns of interoceptive and exteroceptive attention to understand if interoceptive attention is a distinct process;
2. Further distinguish between the neural patterns of interoceptive and exteroceptive attention using more stringent tests to determine the robustness of the models;
3. Decode moment-to-moment interoceptive attention during sustained attention to explore interoceptive training effects; and
4. Explore the relationship between “objective” interoceptive attention decoded by machine learning models based on neural patterns and “subjective” interoceptive awareness and affective distress based on self- and observant-reports.

The study featured a 2 (Group: MABT vs. Control) × 2 (Session: Baseline vs. Post-Intervention) randomized control design. The Interoceptive/Exteroceptive Attention Task (IEAT) used for classification analysis focused on four conditions: Active Interoception and Exteroception, which involved continuous button-press tracking of respiration and a visual stimulus respectively, and Passive Interoception and Exteroception, which involved passive monitoring of respiration and visual targets in the absence of behavioral tracking.

The clinical trial was registered with ClinicalTrials.gov (NCT03583060) and pre-registered with the Open Science Framework (OSF; https://osf.io/y34ja). All study materials and code are available on the OSF (https://osf.io/ctqrh/). Univariate analysis of the IEAT task is described in a separate manuscript that is published as a preprint (https://biorxiv.org/cgi/content/short/2022.05.27.493743v1), including quality control analyses and demonstrations of equivalent difficulty between task conditions. This study was reviewed and approved by the institutional review board at the University of Washington in accord with the World Medical Association Declaration of Helsinki.

## Results

### Aim 1: Distinguishing Between Within-Session Neural Patterns of Interoceptive vs. Exteroceptive Attention

The first aim of the study was to evaluate whether machine learning classifiers could distinguish between neural patterns associated with four experimental conditions: [Active vs. Passive] monitoring of [Interoceptive vs. Exteroceptive] attention. First, we tested a two-state model contrasting both Interoception conditions against both Exteroception conditions. Second, we evaluated a four-state model featuring: Active Interoception, Passive Interoception, Active Exteroception, vs. Passive Exteroception. Finally, we applied a two-state model again, focusing on the distinction between Active Interoception and Active Exteroception. As a liberal test of discriminability, classification was first performed *within-session*, i.e., trained and tested using data from the same assessment session at either baseline or post-intervention.

#### Within-Session Classification: Interoceptive vs. Exteroceptive Attention

The first model classified all interoception trials against all exteroception trials, collapsing together active and passive conditions. The classifier achieved a 73% accuracy at baseline and 72% accuracy at post-intervention (**Figure 1A**). Participants at both baseline and post-intervention were all classified with an accuracy > 55.4%, the *p* < .05 threshold for chance classification except for one participant at post-intervention who was slightly below chance. However, inspection of individual participant classification revealed considerable heterogeneity between well-classified and poorly classified participants (**Figure 2**). Inspection of the confusion matrices indicated that model was not biased in its errors, i.e., it did not make any systematic errors in predicting one condition than the other (**Figure 1B**).

**Figure 1.**
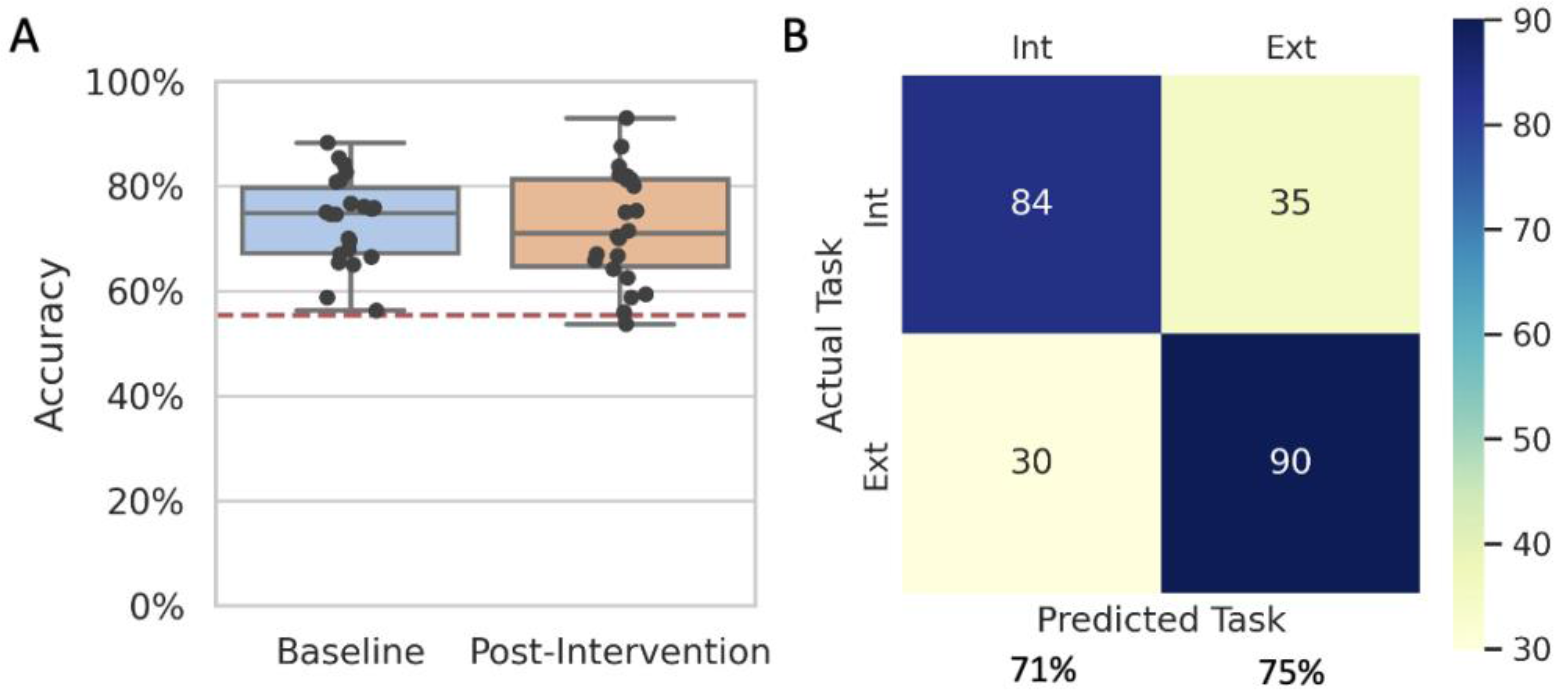
Within-session IEAT classification accuracy: Interoception vs. Exteroception. **(A)** Classification accuracy scores between interoceptive and exteroceptive attention at baseline (73% accuracy) were replicated at post-intervention (72% accuracy). The box represents the quartiles of the dataset; the whiskers extend to show the rest of the distribution, except for data points beyond 1.5 times of the interquartile range which were considered outliers. The data points represent each participant’s classification accuracy score. The red dotted line represents the threshold for individual participants’ overall classification accuracy to be considered statistically above chance. **(B)** The dark diagonal of this confusion matrix shows that the machine learning models did not make any systematic errors. Refer to the **Supplementary Materials** for individual participants’ confusion matrices.

**Figure 2.**
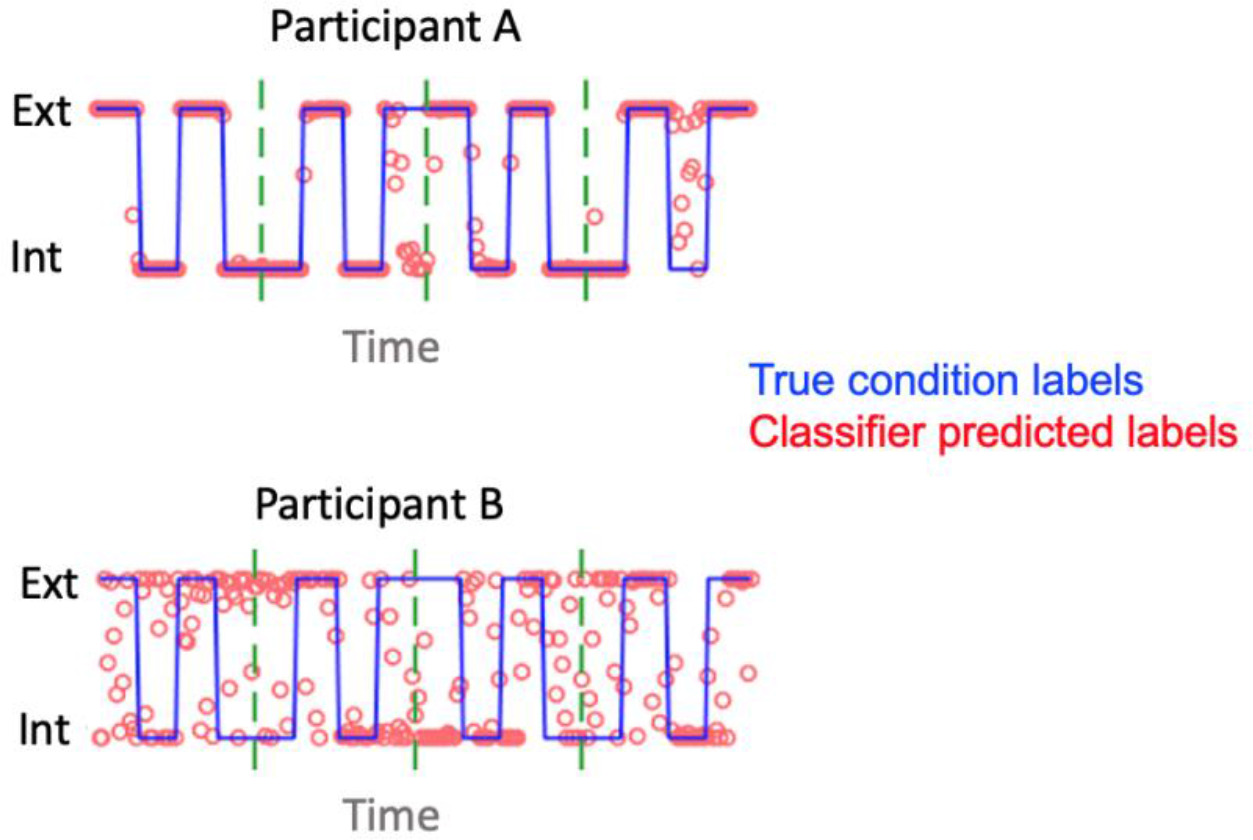
Sample IEAT classification output. The red circles designate classifier predicted labels; The blue lines designate actual condition labels. Participant A’s machine learning model predicted the correct labels over 90% of the time, as can be seen by most of the red circles coinciding with the blue line. Participant B’s model was less accurate, as a large portion of the red circles did not coincide with the blue line.

We generated a group-level frequency map to examine the voxels that contributed the most to the classification (**Figure 3**). Overall, 89% of the important voxels were only important for 2 or fewer participants (**Supplementary Figure 3**); no voxel was important for more than 14 participants. The posterior cingulate and middle insula both contributed evidence for interoceptive attention, whereas the ventromedial prefrontal cortex (vmPFC), dorsomedial prefrontal cortex (dmPFC), motor and somatosensory areas, the primary visual cortex (V1), and the middle temporal visual area (V5) all contributed evidence for exteroceptive attention.

**Figure 3.**
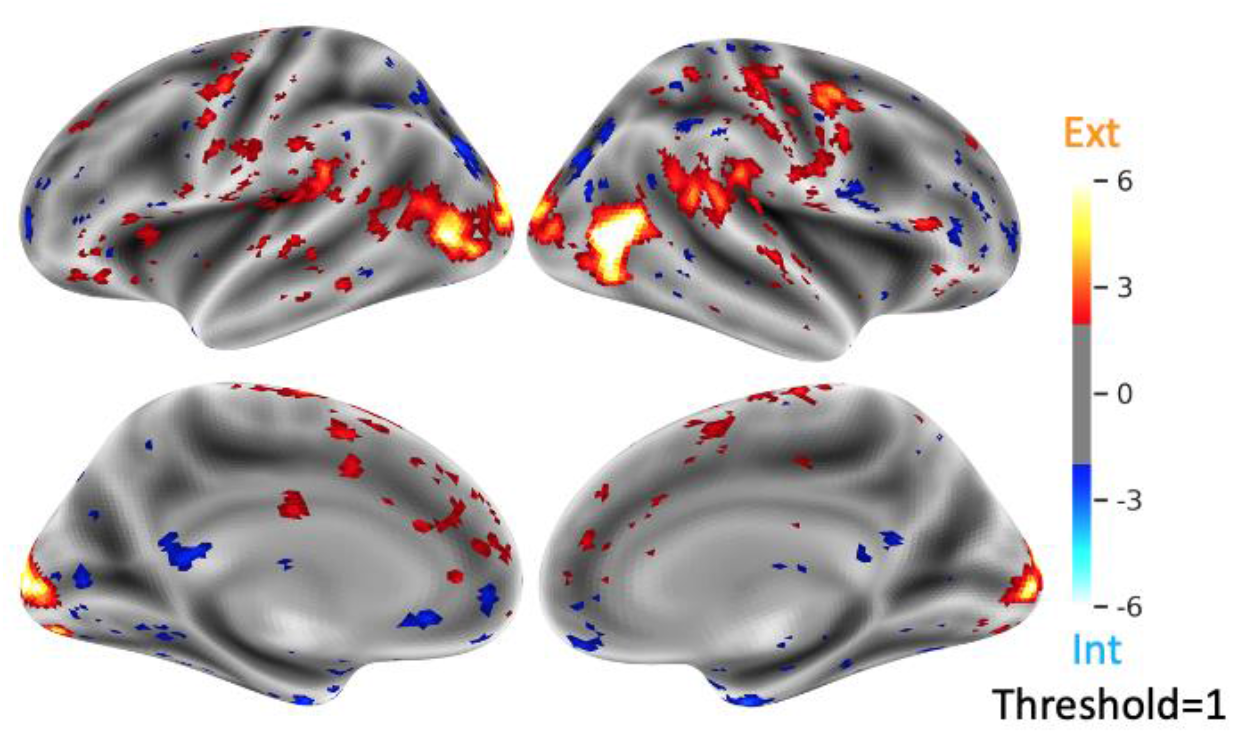
Important voxels in the classification of interoceptive and exteroceptive attention. Regions in red contributed to the classification of exteroceptive attention; regions in blue contributed to the classification of interoceptive attention. These maps were aggregated across all participants; the visualization was thresholded to show voxels that were important to more than one participant. Top panels are left and right lateral views; bottom panels are left and right medial views.

#### Within-Session Classification: Active Interoception, Active Exteroception, Passive Interoception, vs. Passive Exteroception

We then examined classification performance among a more nuanced model that aimed to distinguish between the four experimental conditions. The whole-brain L2 regularized logistic regression classifier achieved a 71% accuracy at baseline and at post-intervention (**Figure 4**). In addition, each mental state was differentiated from the other three states with above chance accuracy: Active Interoception = 72%, Active Exteroception = 73%, Passive Interoception = 69%, and PastExtero = 70% (**Figure 4**). Participants at both baseline and post-intervention were all classified with an accuracy > 29.6%, the *p* < .05 threshold for chance classification. Inspection of the confusion matrices indicated that model was not biased in its errors, i.e., it did not make any systematic errors in predicting one condition over the others, although within-passive active and within-passive confusions were numerically greater than confusions between active/passive condition combinations.

**Figure 4.**
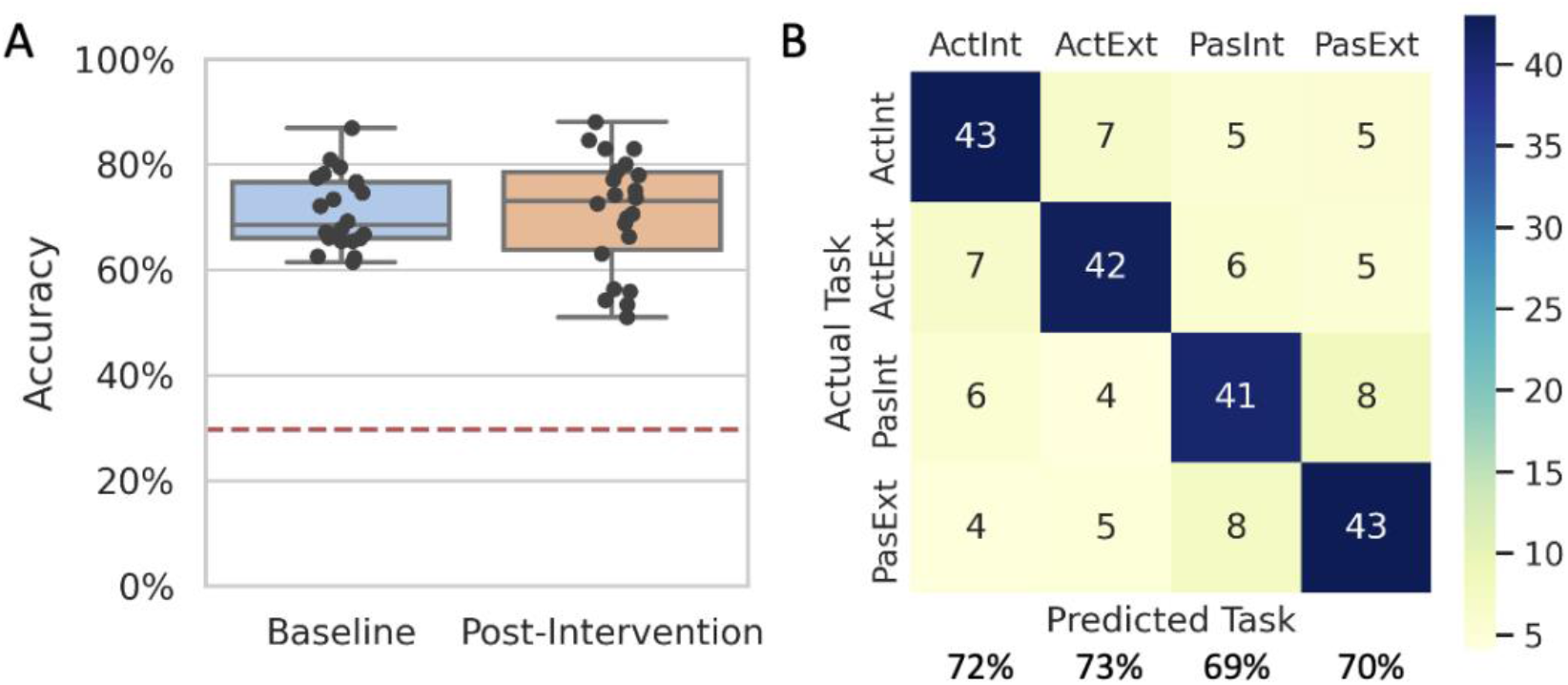
Within-session IEAT classification accuracy: Active Interoception, Active Exteroception, Passive Interoception, vs. Passive Exteroception. **(A)** Classification accuracy scores between the four IEAT conditions at baseline (71% accuracy) were replicated at post-intervention (71% accuracy). **(B)** The dark diagonal of this confusion matrix shows that the machine learning models did not make any systematic errors. Refer to the **Supplementary Figure 2** for individual participants’ confusion matrices.

Next, we generated group-level frequency maps to examine the voxels that contributed the most to the classification (**Figure 5**). Overall, 90% of the important voxels were important for only 2 or fewer participants (**Supplementary Figure 3**); no voxel was important for more than 12 participants in the classification of Active Interoception, 18 participants in the classification of Active Exteroception, 19 participants in the classification of Passive Interoception, and 15 participants in the classification of Passive Exteroception.

**Figure 5.**
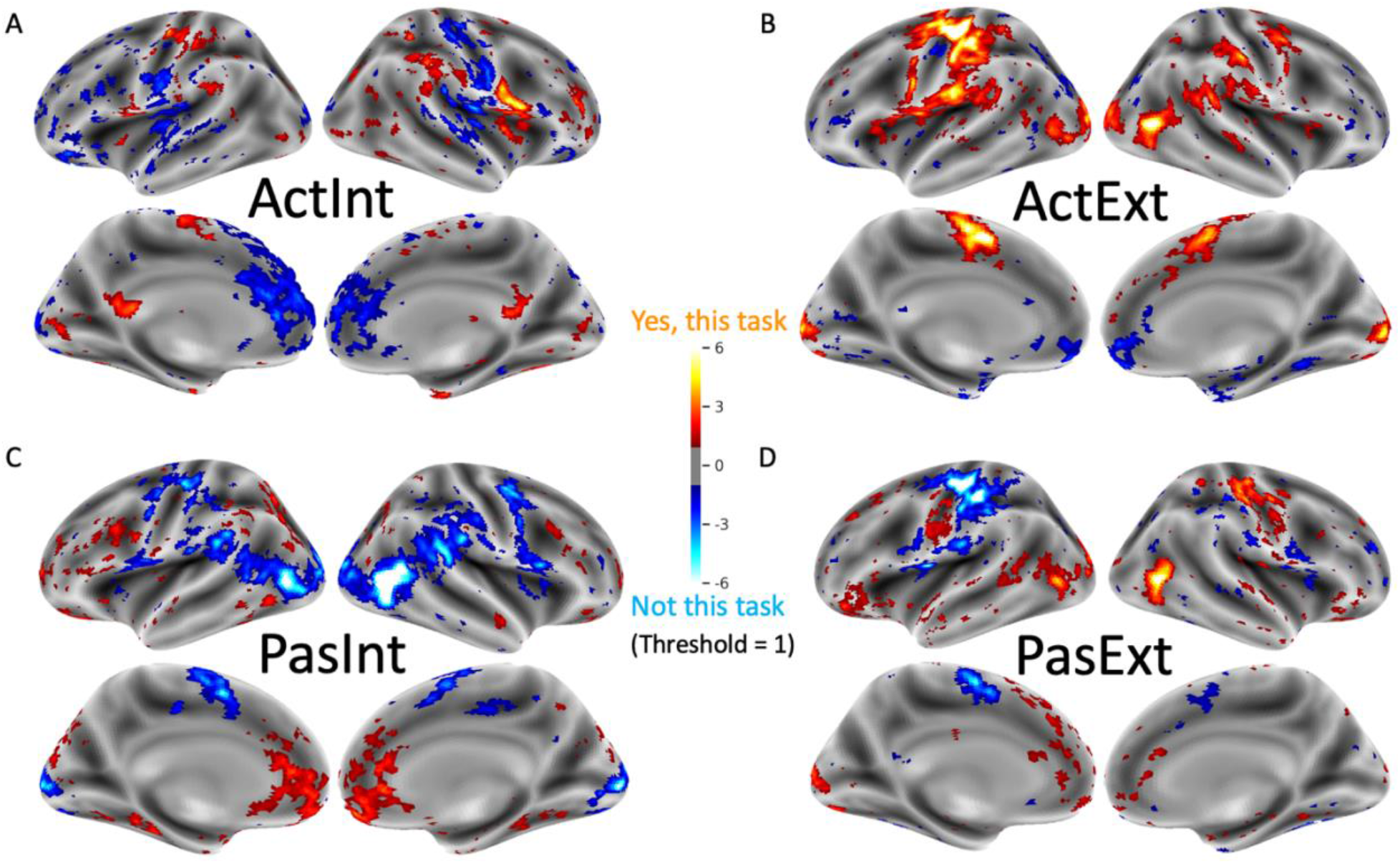
Important voxels in the classification of Active Interoception, Active Exteroception, Passive Interoception, vs. Passive Exteroception. Regions in red contributed to the classification of a specific task (**A**: Active Interoception, **B**: Active Exteroception, **C**: Passive Interoception, **D**: Passive Exteroception). Regions in blue contributed to the classification against that task. These maps were aggregated across all participants; the visualization was thresholded to show voxels that were important to more than one participant.

In keeping with the prior literature, the posterior cingulate and middle insula both contributed evidence for Active Interoception, whereas the vmPFC, dmPFC, and primary somatosensory cortex contributed evidence against Active Interoception. The ventral visual pathway, especially V1 and V5, contributed evidence for Active Exteroception; a cluster in the medial orbital prefrontal cortex (moPFC) contributed evidence against Active Exteroception. The perigenual anterior cingulate cortex (pACC), vmPFC, and moPFC contributed evidence for the Passive Interoception condition; the ventral visual pathway, especially V1 and V5, and the medial premotor cortex all contributed evidence against the Passive Interoception condition. Lastly, V1, V5, the right motor and somatosensory areas, and some small clusters in the prefrontal cortex contribute evidence for Passive Exteroception condition; the medial premotor cortex, left motor cortex, and left somatosensory cortex contributed evidence against Passive Exteroception.

However, in addition to regions supporting representation of interoceptive and exteroceptive content, some classification seemed to capitalize on unequal reporting demands across conditions: for example, motor and somatosensory areas corresponding to button presses with the right hand contributed significantly to Active Interoception and Active Exteroception classification. As mentioned above, area V5 contributed to classification of Passive Interoception, the only condition which did not feature motion. In response, an additional two-state classification analysis was conducted, focused on the most closely matched conditions, Active Interoception vs. Active Exteroception.

#### Within-Session Classification: Active Interoception vs. Active Exteroception

Whole-brain classification between the closely matched Active Interoception and Active Exteroception conditions achieved an 85% accuracy at baseline and 82% accuracy at post-intervention across participants (**Figure 6**). Once again, participants at both baseline and post-intervention were all classified with an accuracy > 55.4%, the *p* < .05 threshold for chance classification. Inspection of the confusion matrices indicated that model was not biased in its errors, i.e., it did not make any systematic errors in predicting one condition than the other.

**Figure 6.**
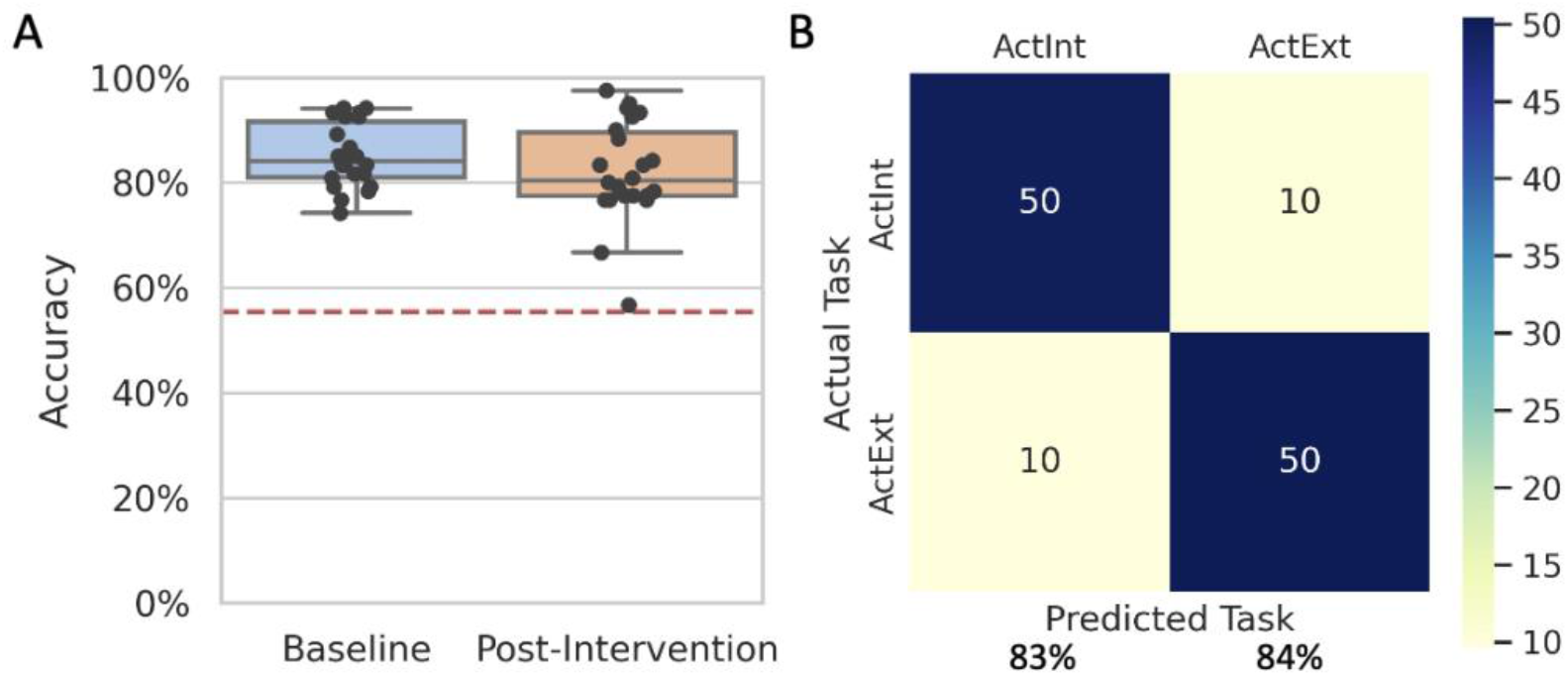
Within-session IEAT classification accuracy: Active Interoception vs. Active Exteroception. **(A)** Classification accuracy scores between the Active Interoception and Active Exteroception conditions at baseline (85% accuracy) were replicated at post-intervention (82% accuracy). The dotted red line represents the threshold for the accuracy scores to be significantly different than chance based on the binomial probability distribution. **(B)** The dark diagonal of this confusion matrix shows that the machine learning models did not make any systematic errors. Refer to the **Supplementary Materials** for individual participants’ confusion matrices.

We generated a group-level frequency map to examine the voxels that contributed the most to the classification (**Figure 7**). Overall, 90% of the important voxels were only important for 2 or fewer participants (**Supplementary Figure 3**); no voxel was important for more than 10 participants. Regions identified in the four-condition classification also demonstrated importance in this analysis. Specifically, the posterior cingulate and middle insula both contributed evidence for Active Interoception, whereas the vmPFC, dmPFC, motor and somatosensory areas, V1, and V5 all contributed evidence for the Active Exteroception condition. Although these two active tracking conditions were matched for motion and button-press requirements, sensorimotor activity supporting classification of Exteroception in the four-condition model was still retained in this two condition model.

**Figure 7.**
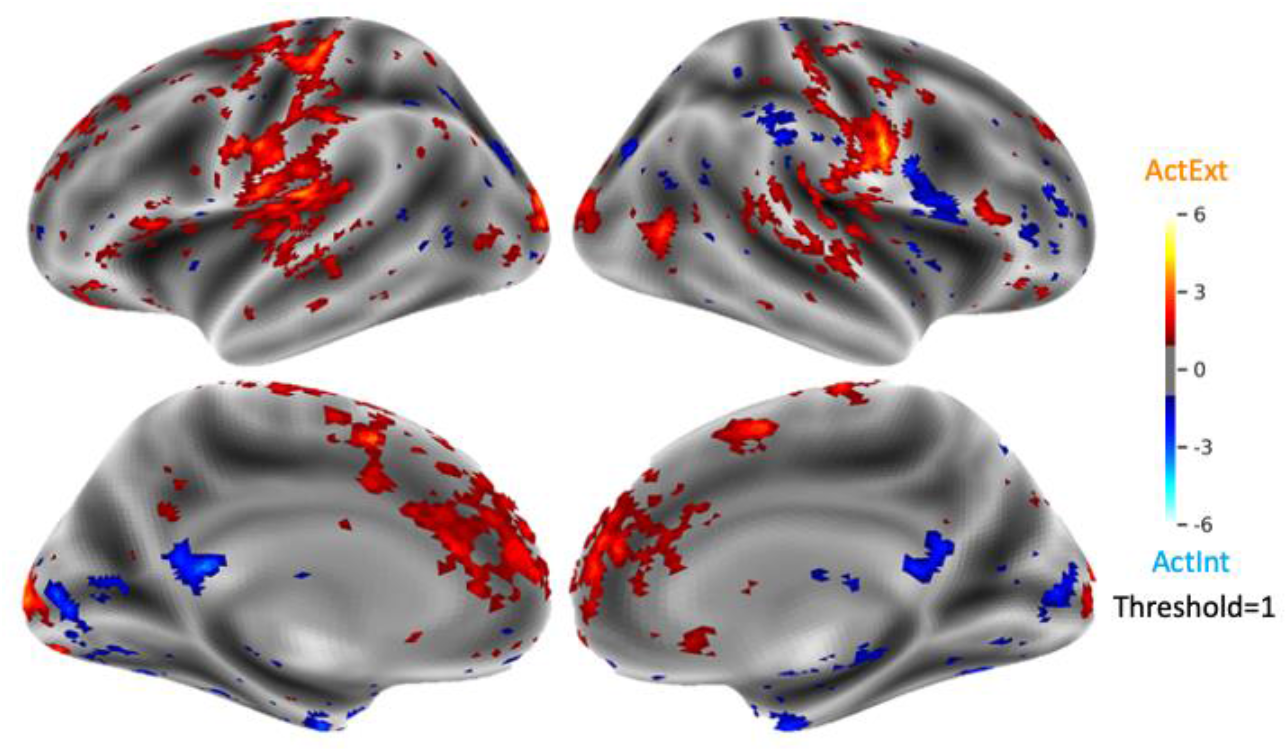
Important voxels in the classification of Active Interoception vs. Active Exteroception. Regions in red contributed to the classification of exteroceptive attention; regions in blue contributed to the classification of interoceptive attention. These maps were aggregated across all participants; the visualization was thresholded to show voxels that were important to more than one participant.

### Aim 2: Distinguishing Between Out-of-Sample Neural Patterns of Interoceptive and Exteroceptive Attention

The second aim of the study was to test whether machine learning classifiers could *predict* individualized neural patterns associated with different attentional states using out-of-sample data. As all participants attended two assessment sessions (baseline and post-intervention) we tested classification models derived from one session on the other session, which was not used to develop that specific model. In other words, models trained on baseline data were tested on post-intervention data and vice versa. We applied this approach for both the four condition (Active Interoception, Active Exteroception, Passive Interoception, and Passive Exteroception) and two condition (Active Interoception and Active Exteroception) models.

The four-condition classifier achieved 51% accuracy across participants when trained on baseline data and tested on post-intervention data, and 50% accuracy when trained on post-intervention data and tested on baseline data (**Figure 8A**). Although less accurate than within-session classification, the overall classification remained significantly above chance.

**Figure 8.**
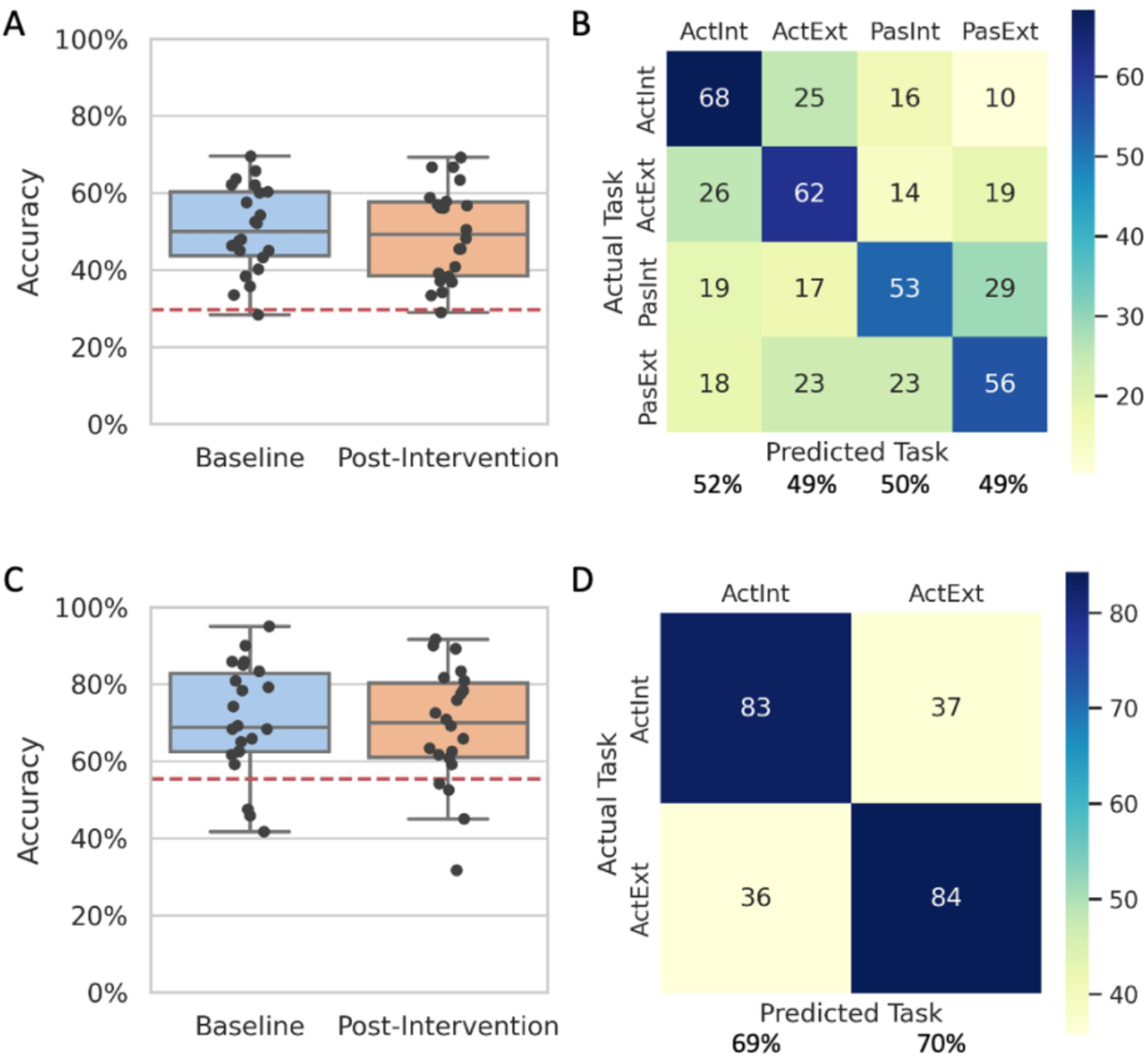
Out-of-sample IEAT classification accuracy. **(A)** Classification accuracy scores between the four IEAT conditions at baseline (51% accuracy) were replicated at post-intervention (50% accuracy). The box shows the quartiles of the dataset; the whiskers extend to show the rest of the distribution, except for data points beyond 1.5 times of the interquartile range which were considered outliers. The data points represent each participant’s classification accuracy score. The dotted red line represents the threshold for the accuracy scores to be significantly different than chance based on the binomial probability distribution. **(C)** The Active Interoception-Active Exteroception classification accuracy scores were 71% at baseline and 69% at post-intervention. **(B) & (D)** Confusion matrices show that the models did not make systematic prediction errors by consistently mistaking one condition for another. Refer to the **Supplementary Figure 4** for individual participants’ confusion matrices.

The binomial probability distribution determined that classification accuracy scores above 29.4% were statistically significant above chance (cumulative probability < .05 for a 25% chance level across all TRs). Of all participants, one failed to surpass chance at baseline (accuracy = 28%) and a different participant failed to surpass chance at post-intervention (accuracy = 29%). Confusion matrices suggested that each state was equally distinguishable from the other three states with above chance accuracy: Active Interoception = 52%, Active Exteroception = 49%, Passive Interoception = 50%, and Passive Exteroception = 49% (**Figure 8B**).

In the two-condition classification model between Active Interoception and Active Exteroception, the whole-brain L2 regularized logistic regression classifier achieved an 51% accuracy at baseline and 50% at post-intervention (**Figure 8C**). The binomial probability distribution determined that classification accuracy scores above 55.4% were significantly above chance (cumulative probability < .05 for a 50% chance level across all TRs). Three participants failed to beat chance at baseline and four at post-intervention. Similar to the four-condition classification, the machine learning models did not make any systematic errors over-estimating one condition than the other: Active Interoception = 69% accuracy and Active Exteroception = 70% accuracy (**Figure 8D**).

### Aim 3: Decoding Interoceptive Attention During Sustained Interoceptive Attention

Our third aim was to explore whether machine learning classifiers trained on the IEAT data could decode participants’ attentional state over multiple three-minute period of sustained interoceptive attention. We were particularly interested in exploring whether interoceptive training influenced these sustained attention metrics.

We first applied the four-condition IEAT classification model to the sustained attention data, which afforded decoding of (i) task demands (active vs. passive attention), (ii) attentional target (interoception vs. exteroception), and (iii) the interaction between task-demands and attentional target. The aim of using the four-condition model was to examine where participants most clearly showed active engagement with the sustained attention task, as this part of the timeseries would be most sensitive to training effects.

In the four-condition analysis, classifiers were trained on each participant’s IEAT data and applied to predict the same participant’s moment-to-moment attention during the SIAT. At each timepoint, the classifier predicted one mental state out of four possible states based on the amount of evidence in favor of each state. **Figure 9** shows the overall estimated frequency for all participants across the four conditions and the trajectory of attentional states during SIAT. Overall, there were more active than passive states. A closer look at the time-course revealed that participants were more frequently decoded as being in Active than Passive states during the first half of the task; active tracking prevalence slowly decreased in the second half of the task to match passive monitoring in frequency **(Table 1)**. Exteroceptive attention was more frequent in the first half of the task, and then decreased over time to match interoceptive attention levels.

**Figure 9.**
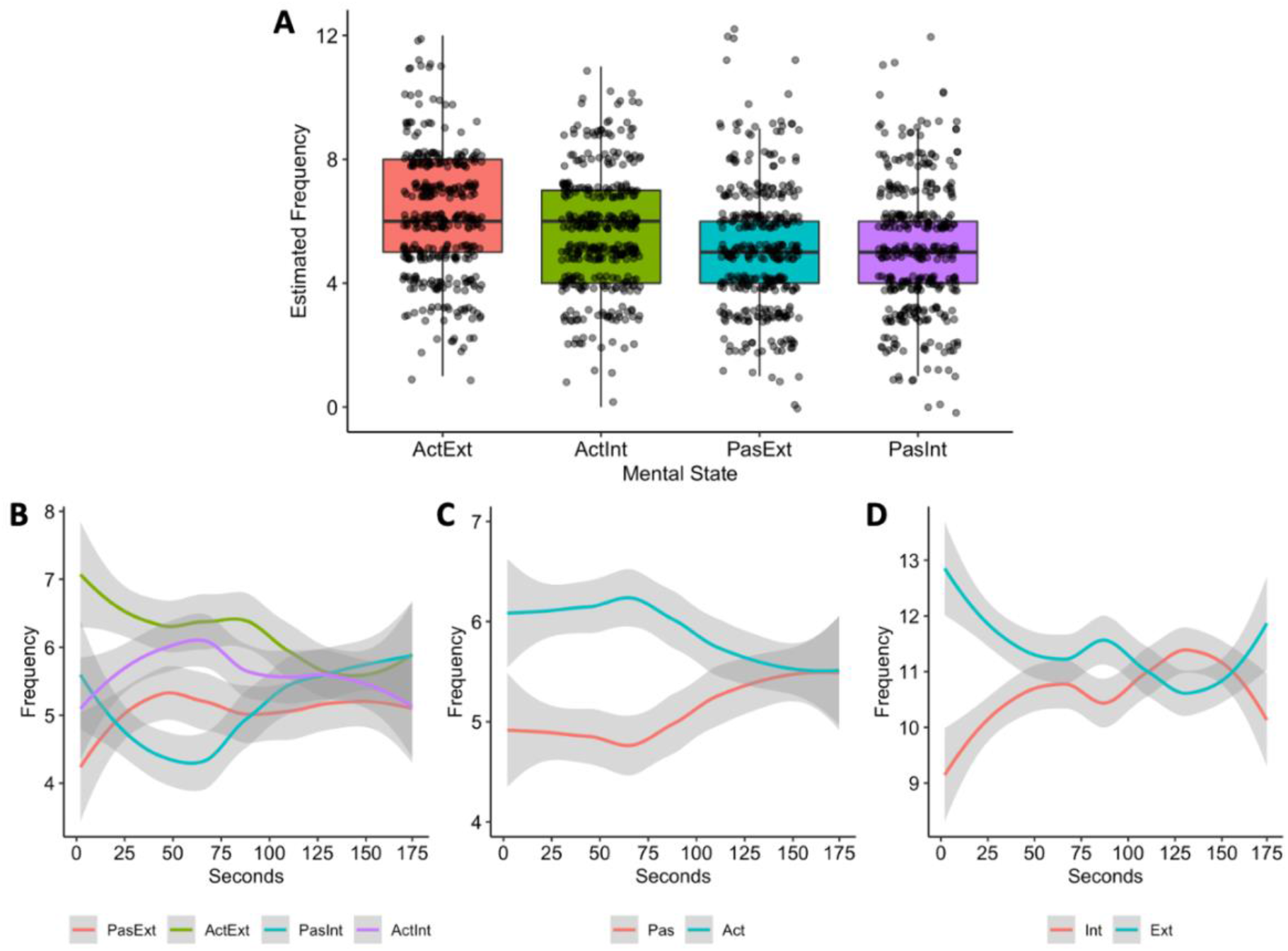
Estimated frequency of mental states during the SIAT. **(A)** The machine learning classifiers predicted more active than passive mental states and more exteroceptive than interoceptive mental states during sustained interoceptive attention. Black dots indicate individual participants’ datapoints. **(B)** The frequency, in terms of the number of participants, of each mental state predicted by the four-condition classifier. **(C)** The frequency of passive and active states over time. **(D)** The frequency of Active Interoception and Active Exteroception states over time.

**Table 1.** Modeling Predicted Frequency of Mental States Using Reporting Demand, Attentional

| *Predictors* | **Predicted Frequency** |  |  |
| --- | --- | --- | --- |
|  | *Estimates* | *CI* | *p* |
| (Intercept) | 19.84 | [18.35; 21.33] | **<0.001** |
| TR | 0.01 | [-0.02; 0.04] | 0.355 |
| Reporting Demand | 7.23 | [5.12; 9.33] | **<0.001** |
| Attentional State | -2.01 | [-4.12; 0.10] | 0.061 |
| TR × Reporting Demand | -0.07 | [-0.11; -0.03] | **0.001** |
| TR × Attentional State | 0.05 | [0.00; 0.09] | **0.030** |
| Reporting Demand × Attentional State | -1.79 | [-4.77; 1.19] | 0.238 |
| TR × Reporting Demand × Attentional Target | -0.00 | [-0.06; 0.06] | 0.943 |
| Observations | 348 |  |  |
| R<sup>2</sup> / R<sup>2</sup> adjusted | 0.250 / 0.235 |  |  |

*Note.* Reporting Demand = active reporting vs. passive watching; Attentional Target = exteroceptive vs. interoception attention. Bolded values indicate $p < .05$ .

Given the absence of task confounds in the two Active conditions, and the apparent dominance of active over passive decoding in the first half of sustained attention runs, we next investigated MABT training effects of attention using a more focused classification model between only the two Active conditions. The readout estimates were plotted for MABT and control participants at baseline and post-intervention (**Figure 10**), and a multilevel model featuring Group, Session, and TR was run on the proportion of Active Interoception classifications made at each TR (**Table 2**). A significant Group × Session × TR interaction indicated that Active Interoception was significantly more frequent in the MABT Post-Intervention group than the other Group × Session combinations. Visual inspection suggested that the interaction was driven by differences in the first half of the time series, consistent with the previous analysis finding that active attentional engagement was highest in this first half.

**Figure 10.**
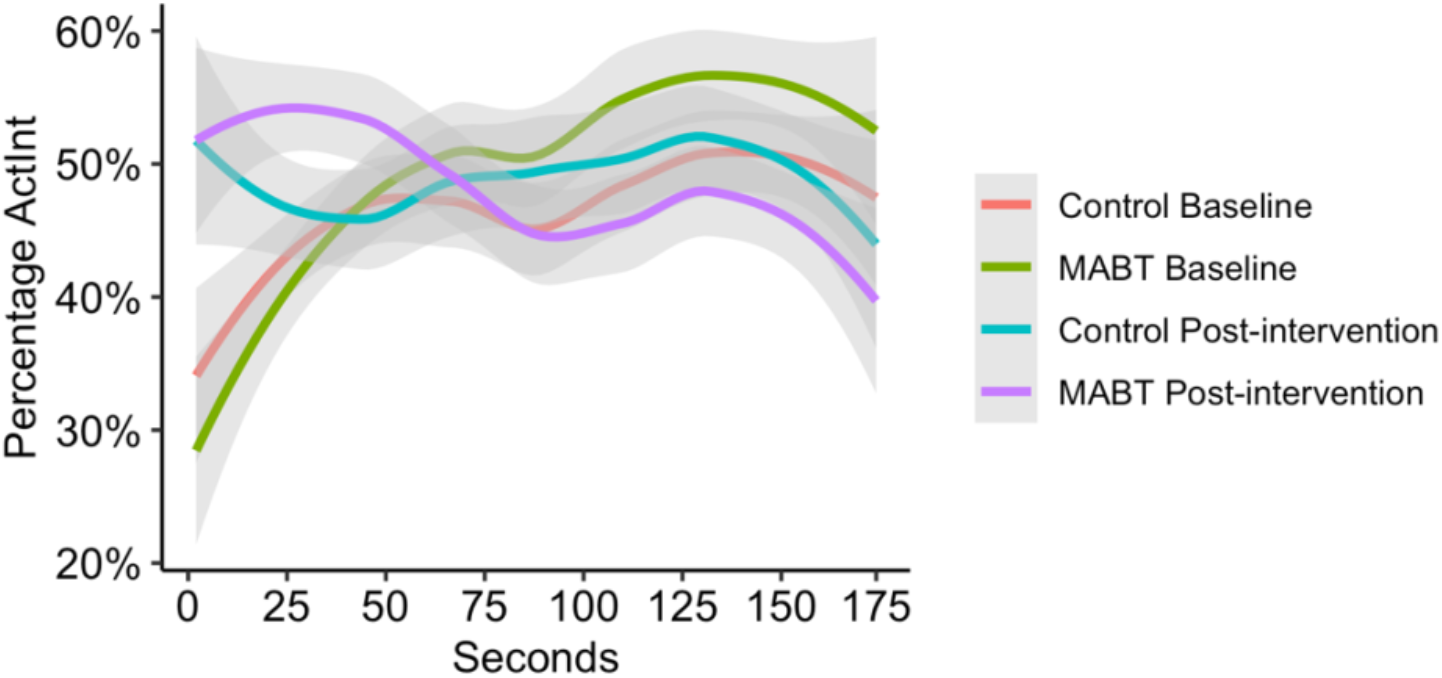
Time-course of estimated percentage of Active Interoception during the SIAT. The moment-by-moment proportion of participants having Active Interoceptive attention (ActInt) in MABT and Control groups at baseline and post-intervention. See **Supplementary Figure 5** for the timecourse of estimated frequency of other attentional states during the SIAT.

**Table 2.** Percentage of Active Interoception in Active Interoception-Active Exteroception classifier estimates of SIAT attention.

| *Predictors* | *Estimat* |  |  |  |
| --- | --- | --- | --- | --- |
|  | *e* | *SE* | *t* | *Pr(>\|t\|)* |
| (Intercept) | 9.14 | 0.45 | 20.41 | **< 0.001** |
| Group | -0.59 | 0.63 | -0.94 | 0.350 |
| Session | 1.21 | 0.63 | 1.91 | 0.057 |
| TR | 0.03 | 0.01 | 3.01 | **0.003** |
| Group × Session | 2.26 | 0.90 | 2.52 | **0.012** |
| Group × TR | 0.03 | 0.01 | 2.25 | **0.025** |
| Session × TR | -0.02 | 0.01 | -1.39 | 0.167 |
| Group × Session × TR | -0.07 | 0.02 | -3.74 | **< 0.001** |

*Note.* Bolded values indicate $p < .05$ . In this two-condition classifier estimate, p-values for Active Interoception and Active Exteroception are identical because the condition was estimated to be either Active Interoception or Active Exteroception with no other solutions.

In an attempted replication of Weng et al (2020), the time-course data was summarized using two derived scores: 1) the total number of events in the SIAT, and 2) the average duration of these events over the course of the SIAT. We used the active-only, two condition classifier (Active Interoception vs. Active Exteroception) to understand these noise-filtered attention estimates. However, statistical models of these new attention metrics using Group, Session, and Attentional Target did not reveal significant training effects (**Supplementary Table 2**). Nevertheless, a marginal finding (*p <* .1) suggested that participants experienced longer duration of events following MABT training (**Supplementary Figure 6**).

### Aim 4: Exploring the Relationship Between Classifier-Estimated Interoceptive Attention and Subjective Interoceptive Awareness and Affective Distress

To better relate the neural classification of interoception to self-reported interoception and wellbeing, we examined the relationship between SIAT estimates based on the two active conditions and MAIA and affective symptom scores using multilevel mixed models. In this analysis, we combined data collected at baseline and post-intervention to increase the inferential power (N = 44). Although the results were not statistically significant (**Supplementary Table 3**), we observed some interesting trends (**Supplementary Figure 7**). It appeared that individuals with a higher MAIA score had longer interoceptive than exteroceptive events compared to those with a lower MAIA score, β = .22; 95% CI [-.11; .56]. For symptom rating, the strongest trend was a main effect of symptom rating, which predicted shorter interoceptive and exteroceptive events alike, β = -.87; 95% CI [-2.16; .42].

In addition to self-reported interoceptive awareness and wellbeing, we explored the relationship between therapist-reported interoceptive awareness and sustained attention readout in the MABT group. We used therapist ratings of participant interoceptive awareness in MABT sessions 5-8 to estimate the average number of interoceptive and exteroceptive events. We did not find any statistically significant effects (**Supplementary Table 4**), but the overall trend suggested that participants with higher therapist ratings of interoceptive attention had fewer interoceptive and exteroceptive events alike, β = -.74; 95% CI [-1.71; 0.23]. (**Supplementary Figure 8**). For duration, higher ratings showed a trend for longer duration interoceptive events, while exteroceptive event duration tended to decrease, β = 1.85; 95% CI [-0.84; 4.53].

## Discussion

The present study successfully demonstrated that machine learning classifiers can differentiate between neural patterns of interoceptive attention and exteroceptive attention. Following within-sample classification, we demonstrated true prediction by using a more stringent out-of-sample test, predicting each participants’ attentional state on test data that was independent from the training set, and acquired more than eight weeks apart. Together, these findings suggest that whole-brain fMRI does provide sufficient information to distinguish between interoceptive and exteroception attention.

We then modelled how classification of task data could be applied to decode participants’ dynamic attentional states during a sustained attention task. We were then able to leverage the clinical intervention context in which the study was conducted, demonstrating sensitivity to training effects on the duration of time participants were able to sustain an active interoceptive state. Finally, we were able to illustrate how such decoding could also be applied to continuous covariates such as subjective interoceptive awareness and affective symptom burden. While the results of the decoding analysis were rarely statistically significant, and thus would be inappropriate for inference to the broader population, the analysis results did suggest a positive link the duration of active interoception and wellbeing.

### Distinguishing Between Interoceptive and Exteroceptive Attention

A series of machine learning models each achieved high levels of accuracy in distinguishing between Interoceptive and Exteroceptive attention, in both in-sample and more stringent out-of-sample tests. Greater posterior cingulate and middle insula activity provided evidence for interoceptive attention, in accordance with the literature on these areas’ involvement in interoceptive processing (Craig, 2003; Farb et al., 2013). Conversely, greater cortical midline activity consistent with the brain’s Default Mode Network contributed evidence against interoceptive attention, which could suggest a break away from effortful, elaborative cognitive processing during internal body focus (Raichle & Snyder, 2007). It is important to note that very few voxels were important for more than two participants in classifying different mental states; this overlap is insufficient for us to identify these regions as ‘shared regions’ among participants. For this very reason, machine learning revealed brain regions that may be missed by univariate analysis due to the lack of overlap among participants.

The ability to make more nuanced distinctions between active and passive reporting of the sensory targets was limited by non-equivalent task demands and stimulus presentation between the IEAT conditions. The four-category model revealed the opportunistic nature of the machine learning classifier, which capitalized on two design confounds. First, the active and passive conditions differed in their requirement of button-presses to track sensory targets. Accordingly, the four-category classifier relied heavily on left motor cortex activity to distinguish active from passive monitoring, consistent with the active monitoring requirement to perform right hand finger movements (e.g., Lotze et al., 2000; Rao et al., 1995). Second, Passive Interoception was the only condition in which the visual target (the circle stimulus) remained stationary on the screen rather than following a cycle of expansion and contraction. Accordingly, the classifiers relied on the differential neural representations in the visual pathway, especially V1 and V5, to distinguish Passive Interoception from the other three conditions, consistent with the literature on these visual areas’ function in mapping the visual field and coding for motion (see Greenlee & Tse, 2008 for a review). This reliance of the classifier on task difference highlights the need to develop closely matched experimental conditions if one wishes for machine learning algorithms to base classification constructs of interest. Future research might revisit the four-category model if these task confounds can be better addressed.

Given the fixation of the general two-category and four-category classification models on nuisance covariates, we refocused our analyses only on the two active conditions (Active Exteroception and Active Interoception). The Active Tracking provided empirical data that participants were engaged and had equivalently facility for performing the two active tracking tasks (see the related preprint https://biorxiv.org/cgi/content/short/2022.05.27.493743v1 for analyses confirming equivalent tracking performance between the two tasks). As mentioned above, the two Active Tracking conditions were also well-matched for stimulus dynamics, and both required button-press tracking of the appropriate sensory target (visual circle or respiratory cycle).

The results supported the feasibility of classifying between carefully controlled periods of interoception and exteroception. Even without the evidence provided by motion and motor confounds, the classification accuracy between Active Interoception and Active Exteroception was the highest of the three models tested, surpassing a more general model that collapsed together active and passive conditions, as well as the four-category model. The classification revealed a similar set of brain regions as the previous four-condition analysis, excluding the motor and visual cortex confounds described above. What remained was a consistent account of the neural distinction between interoception and exteroception: the posterior cingulate and middle insula contributed evidence for Active Interoception, consistent with previous studies that identified these areas as being implicated in the interoceptive neural network (Li et al., 2017; Matsumoto et al., 2006). For Active Exteroception, areas including the vmPFC, dmPFC, motor and somatosensory areas, as well as the visual cortices contributed important evidence.

Why was exteroceptive attention associated with greater reliance on non-visual brain areas despite having the same visual stimulus as the interoceptive condition? One possible explanation is that greater attentional resources might have been required for Active Exteroception to track and report the movements of the visual stimulus as part of the task requirement. Conversely, Active Interoception recruited fewer resources in the cortical midline structures which are associated with cognitively demanding tasks (Seeley et al., 2007). Interoceptive attention might have shifted cognitive resources from elaborative cognitive processing to an inward, sensory focus as suggested by the engagement of the cingulate cortex and insula (Farb et al., 2007, 2010, 2013).

Another important finding was the maintenance of above-chance classification when classifiers were applied out-of-sample. Classifiers trained on a participant at one time point were still able to classify attentional states using data acquired at a second timepoint, one more than eight weeks from the acquisition of classifier training data. While accuracy predictably declined from within-sample classification, classification remained significantly above chance: approximately 50% accuracy vs. a 25% chance in the 4-state classification, and 70% accuracy vs. 50% chance in the 2-state classification, comparable to similar studies that have attempted to classify between mental states. For example, Mitchell et al., (2003)’s Gaussian Naïve Bayes models achieved approximately 80% accuracy in classifying cognitive states such as reading a word about people vs. buildings (50% chance accuracy). Weng et al. (2020)’s L2 regularization models achieved approximately 41% accuracy vs. 20% chance for distinguishing between five mental states, including interoceptive sensations of breath, sensations of the feet, mind wandering, self-referential processing, and ambient sound listening.

In our study, classification rates decreased by 15% when we classified participants’ data out-of-sample compared to within-sample. However, it is unclear whether this drop in accuracy was a “bug or a feature”: if within-sample classification led to overfitting, the lower out-of-sample classification rates would represent a more accurate and generalizable estimate of model prediction. However, accuracy could also decline because individuals’ neural dynamics truly changed, both because of natural variation over a two-month interval and because half of the participants were engaged in interoceptive training over this period. In this case, greater model inaccuracy between time points could be a sign of the model’s sensitivity to training effects: participants who experienced greater training-related change in the neural dynamics of interoception would presumably show poorer model fit. In our sample, this is not the case as we did not observe any difference in out-of-sample classification accuracy between the training and the control groups. Future research that compares the fidelity of both types of models (within- and out-of-sample) against participant experience or performance could adjudicate between these interpretations.

At present, we can at least conclude that the classification models were somewhat robust against overfitting and still performed well above chance on datasets collected two months apart. The primary aim of the study was therefore successful: whole-brain fMRI of participants toggling between interoceptive and exteroceptive attention produces sufficient information to distinguish between neural modes of interoceptive and exteroceptive attention. While classification accuracy can be improved, there seems to be little doubt that active engagement in interoception and exteroception are neurally dissociable states. Furthermore, while activity in primary representation cortices for interoception and vision were important for the classification, less prefrontal and sensorimotor activity appears was indicative of interoception. Why the interoceptive state seems to involve less cortical activity is a ripe topic for further investigation; however, this finding is consistent with the characterization of interoceptive attention as a reducing unnecessary energy demands and driving hemostatic regulation (Quigley et al., 2021), supporting the conditions for embodied self-awareness as a “minimal phenomenal self” (Limanowski & Blankenburg, 2013) or “proto-self” (Bosse et al., 2008).

### Decoding Sustained Interoceptive Attention

The machine learning models trained on IEAT data were applied to decode three-minute runs of the Sustained Interoceptive Attention Task (SIAT). The general properties of the task were decoded in terms of the task demand [active vs. passive] and attentional target [interoception vs. exteroception] categories. In general, a greater proportion of the SIAT period was decoded as being in an active tracking than passive monitoring state. Two further interactions with were observed. First, active tracking was most prominent early in the sustained attention period, but this advantage diminished over time. This finding is not surprising, given the difficulty inherent to sustaining attention over time-attention should be most focused immediately following the guided audio meditation that preceded each SIAT run, but then deteriorate over time as fatigue and/or distraction sets in. Second, participants tended to be engaged in exteroception more than interoception during these earlier phases. This may be a result of two salient exteroceptive cues - the guided meditation recording, and then the onset of the scanner functional recording, both of which may bias attention away from the intended interoceptive target at the start of each run.

Together, these results suggest some challenges in examining sustained interoceptive attention in the scanner. Attention may be inherently biased towards the noise of the scanner environment, and attention may also cease to be consistently engaged about halfway through each three-minute fMRI scanner run. These considerations provided an important context for our next analyses, suggesting that future research on sustained interoceptive attention might focus most fruitfully on the first few minutes of attention to maximize assessment during periods of participant engagement.

#### Training Effects

Following general characterization of the SIAT, we then examined the influence of MABT intervention training effects upon the neural decoding data. Distinct patterns were observed between the two group, driven by post-intervention differences. At baseline, participants were generally more often engaged in exteroception than interoception. However, at post-intervention MABT participants were showed greater interoception than exteroception, a shift not shared by the control group. This statistically significant training effect therefore qualified the general finding of greater exteroception at the start of each SIAT run. A combination of audio recordings prior to the run, combined with the noise of the fMRI data acquisition, may have created a general bias towards exteroception, but MABT appeared sufficient to overcome this bias and allow for sustainable interoceptive processing. It should be noted that the initial ability to engage interoception in the MABT post-intervention group was still limited by the decline in active engagement over the three-minute run-as active engagement declined, so too did the distinction between the MABT and Control groups.

To explore the data for training effects more fully, we examined several other metrics of interoceptive capacity. First, we defined “stable” periods of attention as 6 consecutive seconds or at the same attention state, filtering out frequent switches between interoceptive and exteroceptive attention which might be driven by noise in the data. We found some preliminary (but statistically insignificant) indications that MABT participants might have experienced more stable periods of interoceptive and exteroceptive attention after the training, suggesting an enhanced ability to sustain attention despite not meeting statistical significance. As discussed earlier, the unfiltered time-course analysis showed significant results, whereas the noise-filtered summary metric did not replicate this pattern. The main difference between these analyses is that the noise-filtered analysis reduced hundreds of TRs into a single summary score which resulted in lower statistical power. Hence, there appeared to be a tradeoff of lower power from this statistical reduction that attempted to simplify accounts at the expense of the power inherent when using complete data in multilevel models. Modelling raw rather than summary decoding data may therefore continue to be a useful technique for more powerfully interrogating participant mental states in future research.

### Decoded Sustained Interoceptive Attention and Perceived Awareness

As an exploratory analysis, we studied the relationship between classifier model-estimated interoception during sustained attention and self-reports of interoceptive awareness and distress. Although the analyses were exploratory and the results were not statistically significant (**Supplementary Table 3**), we observed several interesting trends (**Supplementary Figure 6**): individuals with higher subjective interoceptive awareness appeared to have more stable periods of classifier-estimated interoceptive attention, as did those with less self-reported affective symptom burden. Our exploratory analysis of therapist ratings for participants’ interoceptive awareness did not reveal statistical significance (**Supplementary Table 4**) but showed some interesting preliminary trends as well (**Supplementary Figure 7**): individuals who were believed to have higher interoceptive awareness had slightly fewer switches between exteroceptive and interoceptive attention and appeared to have sustained longer attention on interoceptive than exteroceptive signals.

Based upon these preliminary trends, we speculate that self- and therapist-perceived interoceptive awareness might be reflected at the neural level as more stable (as opposed to more frequent) periods of interoceptive attention. It is also possible that distressed individuals may have a reduced capacity to sustain interoceptive attention, as detected by our fMRI-based classifier models, due to repetitive negative thinking that characterizes affective distress (Gustavson et al., 2018). Future studies need to rigorously test these ideas to advance our understanding of whether and how we can improve wellbeing by enhancing interoceptive abilities.

### Limitations and Constraints on Generality

There were some limitations in the design of the fMRI tasks that can be improved in future iterations of this study. One limitation was the lack of guarantee of participant compliance during the passive conditions of the IEAT in which participants were not required to produce any behavioral responses, making it difficult to verify whether and how much they paid attention to the task. In our analysis, we focused on the active conditions in which participants’ attention was guaranteed through the high accuracy of button presses. Future studies can increase the usability of passive-condition fMRI data, for example, by incorporating occasional catch trials that require button presses.

In this study, we modelled two experimental factors that involved active vs. passive and interoceptive vs. exteroceptive attentional states. Future experiments might consider a greater variety of mental states so that interoception can be further compared to states such as mind-wandering, memory recall, and focused attention on different types of exteroceptive stimuli. Post-scan qualitative interviewing can also be built into future studies to better understand what different attentional processes feel like for the participants at a phenomenological level (Petitmengin, 2021). With more experimental conditions designed, it is critical to keep in mind that these conditions need to be closely matched so that machine learning classifiers can focus on important constructs without leveraging too much on nuisance variables.

Another limitation of this study was that no data were collected during the audio-guided interoceptive practice before the self-guided sustained attention. Since we found significant MABT training effect in the beginning of the self-guided sustained attention during which participants were more engaged in the task, it is likely that a high level of task engagement is needed to scaffold the recently developed interoceptive capacity in the training group. Therefore, future research might benefit from extending data collection to include the guided meditation period or from providing attentional prompts during the self-guided sustained attention.

Individuals with varying degrees of experience in mindfulness or interoceptive trainings might benefit differently from such scaffolded attention, which can be another interesting topic of research. In terms of methods, we trained machine learning classifiers to discriminate between interoceptive and exteroceptive attentional states within individuals rather than across individuals. This analysis allowed us to make predictions and estimates based on each individual’s unique task-related neural signature and generate group-level brain maps to identify common brain areas associated with interoceptive and exteroception. Yet, we have not characterized whether interoceptive attention could be consistently observed across individuals from a single training model, or whether models are differentiated across individuals. Future studies can potentially develop across-individual decoders, with the caveat that such decoding is highly complex, can produce superfluous results, and that interpretations about the decodability of mental states should be made with caution (Jabakhanji et al., 2022).

## Conclusions

In this proof-of-concept study, we showed that machine learning can be a powerful tool to characterize the uniqueness of a mental process as compared to other related processes, in this case, interoceptive versus exteroceptive attention. Since interoception is theorized to be the foundation of our feeling states, meaning in life, and broader appraisals of wellbeing, understanding the neural underpinnings of interoception has huge importance. Machine learning also has tremendous potential in estimating or predicting individuals’ mental states, especially when self-report is not preferrable for reasons such as interrupting the mental process being studied, e.g., sustained attention. Furthermore, machine learning can allow us to examine training-related changes at a neural level, in interoception or other types of mental health interventions. For example, therapies that have elements of interoception can benefit from understanding what role and how big of a role interoception plays in improving wellbeing.

The analyses revealed high accuracy of machine learning models in distinguishing interoceptive from exteroceptive attention and related brain regions. Specifically, the posterior cingulate and middle insula were associated with interoception whereas the somatosensory, motor, and cortical midline regions were associated with exteroception. We then estimated individuals’ moment-to-moment attention when they were instructed to sustain focus on body sensations. We observed promising MABT training effects on interoceptive attention immediately following an audio-guided meditation, showing that individuals new to interoceptive training increased their capacity for interoceptive attention.

Although exploratory, individuals with higher subjective interoceptive awareness and those with lower symptomatic distress appeared to be better at maintaining interoceptive attention. These preliminary findings point to the potential application of machine learning predictions as an objective measure of interoceptive awareness and wellbeing in addition to self-reports. While further research is needed to address the limitations of this study, the present findings showed that interoceptive and exteroceptive attention appeared to recruit distinct neural networks and that machine learning shows promise for advancing our knowledge in interoceptive processes.

## Methods

### Research Protocol

This study was conducted in the context of a 2 (Group: MABT vs. control) × 2 (Session: baseline vs. post-intervention) randomized control trial of MABT to investigate training-related neural changes in the brain. Participants were assessed at baseline before being randomized to the MABT or the control condition and reassessed within four weeks of completing the eight-week MABT intervention period. At both baseline and post-intervention assessments, all participants completed a 20-minute self-report questionnaire that surveyed body awareness and symptoms of distress. Participants then completed a series of fMRI scans, including standard anatomical scans, an Interoceptive/Exteroceptive Attention Task (IEAT), and a Sustained Interoceptive Attention Task (SIAT).

The IEAT consisted of five conditions: Passive Exteroception, Passive Interoception, Active Interoception, Active Exteroception, and Active Matching (**Figure 11**). These conditions varied in terms of Reporting Demand (active reporting vs. passive watching) and Attentional Target (interoceptive vs. exteroceptive attention).

**Figure 11.**
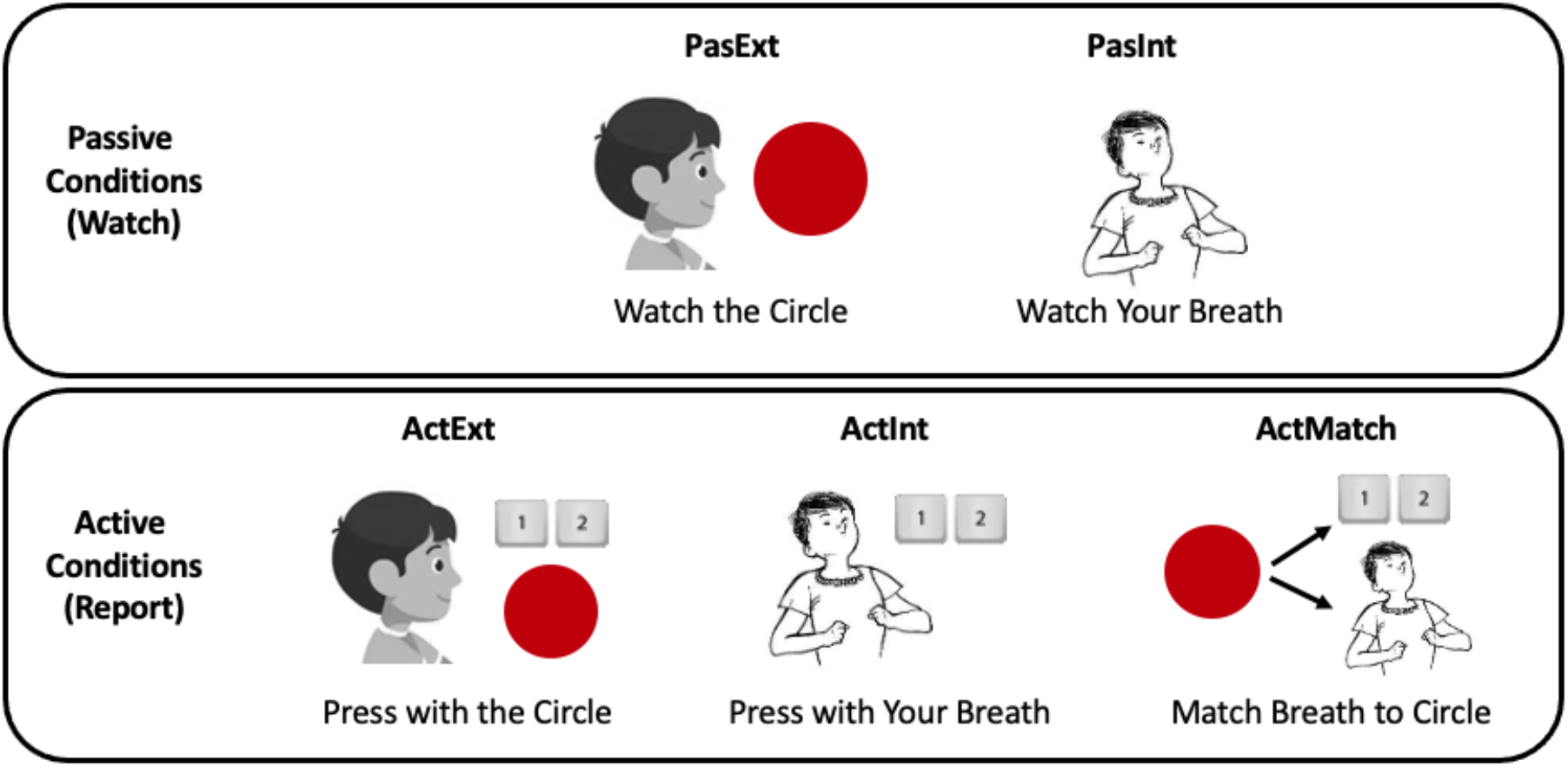
Schematics for the Interoceptive/Exteroceptive Attention Task (IEAT). In the Exteroceptive conditions, participants watched a circle expand and contract; in the Interoceptive conditions, participants paid attention to their inhalation and exhalation. In the Passive conditions, participants simply observed the circle or their breath; in the Active conditions, participants pressed buttons when the circle expanded or contracted and when they inhaled or exhaled. In the Matching condition, participants reported on the circle’s movements while intentionally matching their breathing to the circle’s movements.

### Participants

Participants were recruited through postings on the University of Washington (UW) Institute of Translational Health Sciences website, flyers on campus, as well as advertisements in local newspapers. Advertisements described the study as a mind-body investigation into the neural processes of body awareness for people with moderate stress. Participants provided written informed consent and were compensated for their time. All assessments and data collection took place at the UW Integrated Brain Imaging Center.

Primary inclusion criteria for the study included: (1) over 18 years of age, (2) score on the Perceived Stress Scale meeting screening cut-off for moderate stress, (3) no prior experience with mindfulness-based trainings, (4) agrees to forgo mind-body therapies for the duration of the study, (5) fluent in English, (6) able to attend all study sessions, and (7) right-handed. Primary exclusion criteria included: (1) current diagnosis of any mental health disorder, (2) unable to participate in all sessions, (3) cognitive impairment, (4) head injury involving loss of consciousness, (5) pregnant; or (6) MRI contraindications.

Fifty-seven participants were recruited. Twenty-four participants were excluded for reasons listed in **Supplementary Figure 1**. Twenty-three remaining participants were randomly assigned to receive MABT (n = 12) or no-intervention to serve as the control (n = 11). Out of these participants, 22 completed the required assessments (11 MABT and 11 controls) and were included in the analyses (11 males and 11 females; age range: 18-62 years, mean age = 36.1 years; 20 self-identified as Caucasian, 1 as African American, 2 as Hispanic; highest education: 5 with high school degrees, 2 with 2 years of college, 8 with Bachelor’s degrees, and 7 with Master’s degrees or higher).

Given prior observation of large effects of MABT on subjective interoception (Price, Thompson, Crowell, Pike, et al., 2019), this study was designed to detect medium to large Group × Time interaction effects (f > .3). Power analysis conducted in G*Power software suggested that this sample size would be sufficient to detect such effects with 80% power. This small sample size was sufficient for testing a focal hypothesis anticipating a large training effect in a univariate anlaysis on the same dataset (Farb et al., 2022), but admittedly underpowered to detect weaker downstream effects on clinical symptoms or mechanistic markers.

### MABT

MABT uses an incremental approach to help build comfort and skills needed to develop and facilitate interoceptive awareness (Price & Hooven, 2018). The approach was delivered individually using the manualized 8-session protocol developed for research. This protocol involves three phases: sessions 1-2 focus on body literacy, sessions 3-4 on interoceptive training, and sessions 5-8 on development of sustained mindful interoceptive attention and somatic appraisal processes. A take-home practice is collaboratively developed at the end of each session to facilitate integration of interoceptive awareness in daily life. Eight weekly 75-minute individual sessions were delivered at the UW School of Nursing Clinical Studies Unit by one of two licensed massage therapists trained in the MABT protocol. Protocol compliance was monitored through audio recording of sessions, process evaluation forms, and ongoing clinical supervision. Participants were given up to 10 weeks to complete all 8 sessions to accommodate schedule conflicts. All MABT participants completed at least 75% of the sessions (i.e., at least 6 sessions): Eight participants completed all 8 sessions, two completed 7 sessions, and one completed 6 sessions. Therapists monitored participants’ progress with quantitative ratings and qualitative descriptions on the process evaluation forms completed after each MABT session.

### fMRI Tasks

FMRI data were collected during a novel Interoceptive/Exteroceptive Attention Task (IEAT) and a Sustained Interoceptive Attention Task (SIAT) at both baseline and post-intervention. In each assessment session, there were two fMRI scans (i.e., two runs). Each task was administered once in each functional scan. Altogether, participants performed each task four times in this study, twice at baseline and twice at post-intervention.

#### Interoceptive/Exteroceptive Attention Task (IEAT)

The novel IEAT (Farb et al., 2022) consisted of five conditions: Passive Exteroception, Passive Interoception, Active Interoception, Active Exteroception, and Active Matching as shown in **Figure 11**. These conditions varied in terms of Reporting Demand (active reporting vs. passive watching) and Attentional Target (interoceptive vs. exteroceptive attention).

Each condition started with a 10-second instruction screen followed by a 30-second task period. All conditions were order-counterbalanced and repeated twice in each functional run. Altogether, 6.7 minutes of data were collected in each run and 13.4 minutes in both runs.

##### Passive Conditions

During Passive Exteroception, participants were asked to visually monitor a circle as it expanded and contracted periodically on the MRI-compatible visual display without making any behavioral responses. The circle’s pulse frequency was set to match participants’ estimated in-scanner breathing frequency (usually around 12 Hz). During Passive Interoception, participants viewed a stationary circle on the screen while attending to sensations of the breath.

##### Active Conditions

During Active Interoception, participants were asked to report on their inhalations and exhalations by making key presses with their right-hand index and middle fingers respectively. The circle on the screen also responded to these key presses, approximating the frequency of circle movement during Passive Exteroception. During Active Exteroception, participants were asked to report on the expansion and contraction of the circle on the screen, which again was set to pulse at participants’ in-scanner respiratory frequency.

##### Active Matching Condition

During Active Matching, participants were asked to report on the expansion and contraction of the circle (as in Active Exteroception) by making button presses while matching their inhalation to the circle’s expansion and their exhalation to the circle’s contraction. Together, these five experimental tasks were developed to address the limitations of prior interoception paradigms. However, the goal of the present study was to directly classify between factors of attentional target (interoception vs. exteroception) and reporting demands (active vs. passive monitoring). Because the Active Matching condition required aspects of both interoceptive and exteroceptive attention, it was not used in the classification models that are the focus of this paper, but is reported in a forthcoming univariate analysis paper (https://biorxiv.org/cgi/content/short/2022.05.27.493743v1).

#### Sustained Interoceptive Attention Task (SIAT)

Immediately before fMRI data acquisition, participants listened to a 2.5-minute audio-guided interoceptive awareness meditation in the scanner, directing them to place a hand on their chest and channel mindful attention to the inner space of the chest underneath their hand. After the guided meditation, participants were instructed to sustain attention on inner body awareness for 3 minutes with their eyes closed during fMRI data acquisition. This procedure was repeated across 2 runs at baseline and 2 runs at post-intervention to yield a total of 4 scans, i.e., 12 minutes of fMRI data.

### Questionnaire Measures

During the baseline and the post-intervention sessions, participants answered a self-report questionnaire consisting of the Multidimensional Assessment of Interoceptive Awareness (MAIA), the Patient Health Questionnaire - Somatic, Anxiety and Depressive Symptoms (PHQ-SADS), and the Perceived Stress Scale (PSS). In addition, the MABT therapists rated participants’ capacity for sustained interoceptive attention over the second half of the training period (sessions 5-8). For the descriptive statistics of these self-report measures, see our Open Science Framework page (https://osf.io/ctqrh/).

#### Multidimensional Assessment of Interoceptive Awareness (MAIA)

The MAIA is a 32-item self-report questionnaire used to assess interoceptive body awareness (Mehling et al., 2012). It consists of eight scales each measuring an aspect of interoceptive awareness: Noticing, Not-Distracting, Not-Worrying, Attention Regulation, Emotional Awareness, Self-Regulation, Body Listening, and Trust. These scales have good evidence of internal-consistency reliability with alphas ranging from .66 to .82, and good evidence of construct validity as assessed by inter-scale correlations as well as differential scores between individuals who were expected to have higher or lower body awareness.

#### Composite Affective Symptom Score

A composite affective symptom score was obtained based on responses on the Patient Health Questionnaire - Somatic, Anxiety and Depressive Symptoms (PHQ-SADS; Kroenke et al., 2010) and the Perceived Stress Scale (PSS; Cohen et al., 1983). The PHQ-SADS is a 37-item self-report questionnaire consisting of the PHQ-9 depression scale, PHQ-15 somatic symptom scale, and Generalized Anxiety Disorder (GAD)-7 anxiety scale (Kroenke et al., 2010). All three scales have good evidence of internal-consistency reliability (a = .80 to .92), test-retest reliability (r = .60 to .84), as well as good sensitivity and specificity to detect depression, anxiety, and somatic symptoms. The PSS is a 10-item self-report questionnaire used to assess how feelings and perceived stress levels are affected by various situations (Cohen et al., 1983). A review study showed that the PSS has good internal-consistency reliability (a > .70 in all 12 studies evaluated) and test-retest reliability (r > .70 in all four studies evaluated), although criterion and known-groups validity need to be further evaluated (Lee, 2012). We extracted the first principal component from affective symptom scales PHQ-SADS and PSS, which explained 67.2% of the overall variance. A simulation run using the ’paran’ library confirmed that one factor was sufficient to explain the variances and was used as a composite affective symptom score for this study.

#### Therapist Rating: Capacity for Sustained Interoceptive Attention

In MABT sessions 5-8, therapists rated participants’ capacity for sustained interoceptive attention on a scale of 0-5 based on their observation: 0 = none, 1 = momentary, 2 = fluctuating in and out (being in the state for brief time, i.e., less than 3 minutes), 3 = steady contact for many minutes, 4 = fluctuating in and out (being in the state for longer periods, i.e., more than 3 minutes), 5 = sustained contact (10-30 minutes).

### Data Analysis

#### Imaging Data Acquisition and Preprocessing

Neuroimaging data were collected using a 3T Philips Achieva scanner (Philips Inc., Amsterdam, Netherlands) at the Diagnostic Imaging Sciences Center, University of Washington. Imaging began with the acquisition of a T1-weighted anatomical scan (MPRAGE) to guide normalization of functional images with repetition time (TR) = 7.60 ms, echo time (TE) = 3.52 ms, inversion time (TI) = 1100 ms, acquisition matrix = 256 x 256, flip angle = 7°, shot interval = 2530 ms, and 1mm isotropic voxel size. Functional data were acquired using a T2∗-weighted echo-planar-imaging (EPI) sequence with TR = 2000, TE = 25 ms, flip angle a = 79°, field of view = 240 × 240 × 129 mm, 33 slices, and a voxel size of 3 × 3 × 3.3 mm with 3.3 mm gap. Button presses were registered using a 2-button MR-compatible response pad.

Neuroimaging data preprocessing was performed using the fMRIPrep pipeline 20.0.6 (Esteban et al., 2019) (see **Supplementary Materials** for full details). Preprocessing consisted of realignment and unwarping of functional images, slice timing correction, and motion correction. The functional images were resliced using a voxel size of 2 x 2 x 2 mm and smoothed using a 6-mm FWHM isotropic Gaussian kernel.

#### Analysis Software

The Python Language (Python Software Foundation, https://www.python.org/) was used mainly for machine learning analysis. In-house code was developed with reference to BrainIAK (the Brain Imaging Analysis Kit, http://brainiak.org; (Kumar et al., 2020, 2022). The scikit-learn package was used for machine learning analysis (Pedregosa et al., 2011), the nilearn package for brain maps (Abraham et al., 2014), and the seaborn package for data visualization (Waskom, 2021). The R Language (R Core Team, 2020) was also used for statistical analyses. The lme4 package was used for statistical modeling (Bates et al., 2015) and the ggplot2 package for data visualization (Wickham, 2016).

#### Aim 1: Distinguishing Between Within-Session Neural Patterns of Interoceptive vs. Exteroceptive Attention

We aimed to differentiate neural patterns of interoceptive attention from those of exteroceptive attention and identify brain regions unique to each state. To do so, we trained machine learning classifiers on fMRI data when participants engaged in interoceptive and exteroceptive attention, assessing how accurately these states could be separated and which brain regions contributed to the separation.

Specifically, we used fMRI data collected in the Active Interoception, Active Exteroception, Passive Interoception, and Passive Exteroception conditions. Again, the Active Matching condition was not analyzed in this study, because it blended interoceptive and exteroceptive attention and was therefore deemed unsuitable for distinguishing between these two processes. We conducted the analyses in three steps. First, we combined Active Interoception and Passive Interoception into an interoceptive condition and Active Exteroception and Passive Exteroception into an exteroceptive condition to examine the gross differences regardless of the reporting demand. Secondly, we examined Active Interoception, Active Exteroception, Passive Interoception, and Passive Exteroception as four separate conditions, considering both attentional target and reporting demand. Third, we narrowed in on differences between Active Interoception and Active Exteroception to eliminate the confounds in Passive Interoception, the only condition in which the circle remained stationary on the screen, and to guarantee participants’ engagement by using conditions that required behavioral responses. All three steps were conducted in a similar classification workflow as demonstrated in **Figure 12**.

**Figure 12.**
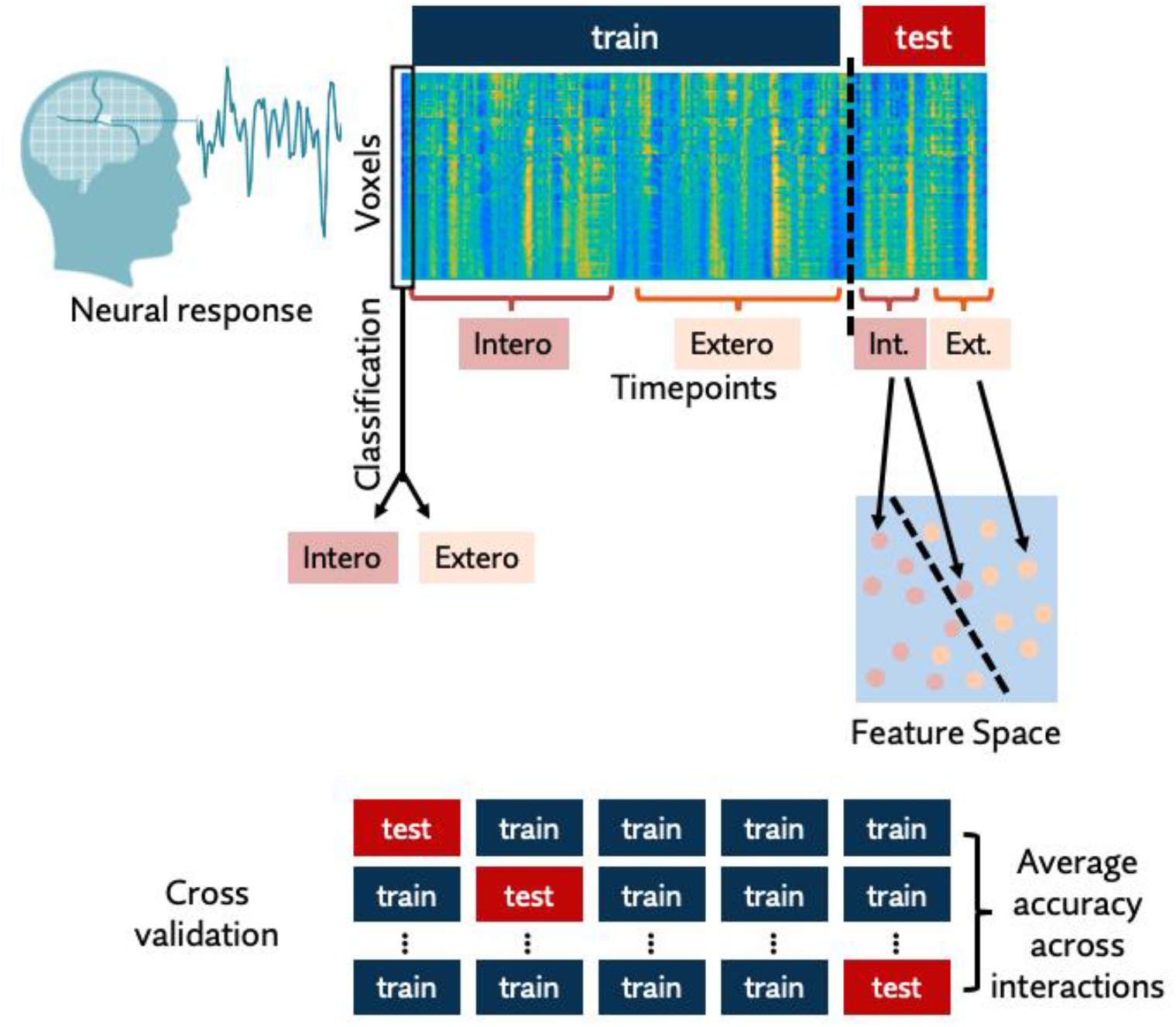
Machine learning classification workflow. First, 4-dimensional fMRI BOLD data were reshaped into a 2-dimensional voxel × timepoints matrix. Then, part of the data was used to train the machine learning model and the other part to test the model. The model used training data to learn weights associated with each voxel and used these weights to predict labels for the test data. Cross validation as performed by assigning different chunks of the dataset as the test set. An accuracy score was calculated by averaging the prediction accuracy across all test sets.

Classification was performed at an individual participant level to maximize classification accuracy and account for individual anatomical and functional differences in the brain. For each participant, the 4-dimensional fMRI data were reshaped into voxel by timepoint matrices. IEAT task-related timepoints were extracted from the full timecourse of the scans. A whole-brain mask was applied to the data to retain voxels that fell within the brain. We used this data-driven whole-brain approach to identify any brain regions that might drive the separation of interoception and exteroception without making *a priori* assumptions about which regions might be critical in the process.

We implemented a penalized logistic regression with L2 regularization (i.e., Ridge Regression) with reference to methods used by Weng et al. (2020) to classify internal states of attention during meditation. Regularization in general penalizes the overfitting of data and prevents the models from over-learning from the training data to the extent that they fail to generalize to out-of-sample data. We selected L2 regularization over other methods such as L1 regularization (i.e., Lasso Regression) because L2 regularization retained more important features (i.e., voxels) and would therefore reveal more brain regions involved in interoceptive and exteroceptive processes. We were aware of many other machine learning algorithms; in our pilot testing, we did compare L2 regularization to methods such as sparse multinomial logistic regression, Gaussian Naïve Bayes, XGBoost, and singular value decomposition linear regression, but failed to see any evidence of superior classification. This lack of distinction therefore led us to proceed with our planned use of the L2 algorithm.

A k-fold cross-validation method was used to train and evaluate the performance of the classifier models. Each participant’s voxel by timepoint matrix in each study session was split into a training set and a test set. In the training set, a logistic regression model used the brain activation values of each voxel at each timepoint as well as the true label of the experimental condition. Each voxel was assigned a weight which indicated how much evidence it provided for or against an experimental condition. Then, the classifier used these weights that it learned from the training set to predict the experimental condition of each timepoint in the novel held-out test set. These predictions were evaluated against the true experimental condition labels to derive a measure of classification accuracy. The classifiers were trained and tested within baseline data and within post-intervention data respectively. This in-sample within-session classification allowed us to evaluate how well the classifiers separated different mental states from the same time of assessment. We split each session’s data into five folds and ran cross validation by training on four folds and testing on the fifth held-out fold (i.e., training on 80% and testing on 20% of the data). This process was repeated until all five folds had been used as the test fold once. An average accuracy score was obtained across the five iterations.

For each participant at each assessment, we applied the binomial theory to analyze whether classification accuracy was significantly greater than chance at the p < .05 threshold. At each session, the four conditions were comprised of 60 volumes each over the two functional runs: (1 volume / 2 sec) * (30 sec / block) (2 blocks / run) * 2 runs * 4 conditions = 240 volumes in total. For the two condition models, the chance probability of successfully classifying a given functional volume was 50% (i.e., choosing one out of two options at random). The binomial probability distribution suggested that participants were each required to achieve classification accuracy above 55.4% to be considered significantly above chance (1 - cumulative probability < .05). For the four condition models, the chance probability was 25%, which by the binomial theorem required classification accuracy above 29.6% to be significantly above chance.

For visualization of the classification models, group level classification maps were created to identify important brain regions. For each participant, voxels that had a major contribution to the classification were identified: Voxels whose weights were above 2 standard deviations of the mean weight were assigned a value of 1, and those whose weights were below 2 standard deviations of the mean weight were assigned a value of −1. Then, all participants’ important maps were overlaid to create a group-level importance map in which higher absolute values indicated more discriminative voxels across participants.

#### Aim 2: Distinguishing Between Out-of-Sample Neural Patterns of Interoceptive vs. Exteroceptive Attention

One step beyond examining within-session classification accuracy was to evaluate whether our models could be generalized across different times of assessment. Specifically, classifiers were trained on the entire baseline data and tested on the entire post-intervention data, and vice versa. Such out-of-sample testing would help us understand whether brain regions critical for the differentiation of interoception and exteroception were stable over time within the same individual.

#### Aim 3: Decoding Interoceptive Attention During Sustained Interoceptive Attention Task (SIAT)

Our next goal was to explore whether we could predict sustained interoceptive attention and examine whether interoceptive training had an impact on how well participants maintained this attention. To do so, we used the classifiers trained on IEAT to estimate participants’ attentional states during SIAT based on their fMRI neural patterns.

First, at an individual level, L2 regularized regression classifiers were trained on each participant’s IEAT data at baseline and post-intervention separately. Then, classifiers trained on baseline IEAT data were used to predict the TR-by-TR labels for the same participant’s SIAT data at baseline, and vice versa for post-intervention. This analysis was conducted primarily on the two active conditions (Active Interoception and Active Exteroception). In addition, comparable results were found when using a four-condition classifier (see **Supplementary Materials** for details).

To characterize participants’ degree of engagement in interoceptive versus exteroceptive attention, a few metrics were developed, including 1) the average amount of time continuously spent in a certain mental state, thresholded at 6 seconds (i.e., 3 TRs) to reduce noise in the prediction (i.e., every continuous period over 6 seconds was considered as an “event”), and 2) the average number of continuous “events” in each mental state. The duration of events and the number of events together offer an estimate of the stability of interoceptive and exteroceptive attention throughout the task.

To explore training effects, multilevel mixed models were used to examine whether MABT improved sustained interoceptive attention. Group membership (i.e., MABT vs. Control) was the between-subjects independent variable; Session (i.e., Baseline vs. Post-Intervention) was the within-subjects independent variable. The frequency or proportion of interoceptive attention as well as metrics for the duration and number of events were used as dependent variables. Significant Group by Session interaction would be regarded as evidence for group-specific training effects.

#### Aim 4: Exploring the Relationship between Classifier Estimated Interoceptive Attention and Subjective Interoceptive Awareness and Affective Symptoms

Our final aim was to explore the relationship between machine learning classifier derived interoceptive attention and subjective perception of interoceptive awareness and wellbeing. Multilevel mixed models were used to predict the duration and number of interoceptive and exteroceptive events using self-reported interoceptive awareness (MAIA) or affective symptom burden. In addition, for the MABT group only, we also examined the relationship between classifier estimates and therapist-reported interoceptive awareness. Multilevel mixed models were used with MAIA, composite symptom scores, and therapist ratings as independent variables and the interoceptive attention metrics, i.e., average duration of events and number of events, as dependent variables.

## Acknowledgements

This work was supported by a RIFP award from the School of Nursing at the University of Washington and a grant from the National Center for Advancing Translational Sciences of the National Institutes of Health [UL1 TR002319]. The content is solely the responsibility of the authors and does not necessarily represent the official views of the National Institutes of Health. Data analysis infrastructure and trainee expenses were supported by a Canadian Natural Sciences and Engineering Council Discovery grant [RGPIN-2015-05901]. We thank Natalie Koh and Sophie Xie for their help in coordination and data collection, and to Dr. Thomas Grabowski, Dr. Chris Gatenby, and Ms. Liza Young at the Integrated Brain Research Center at the University of Washington for their collaboration. We thank Dr. Helen Weng for providing the MATLAB code used for her 2020 paper, which served as a helpful conceptual guide in developing our analysis pipeline. Last but not least, we wish to express our appreciation to the study participants and the MABT therapists, Elizabeth Chaison and Carla Wiechman.

## Supplementary Materials

**Supplementary Figure 1.**
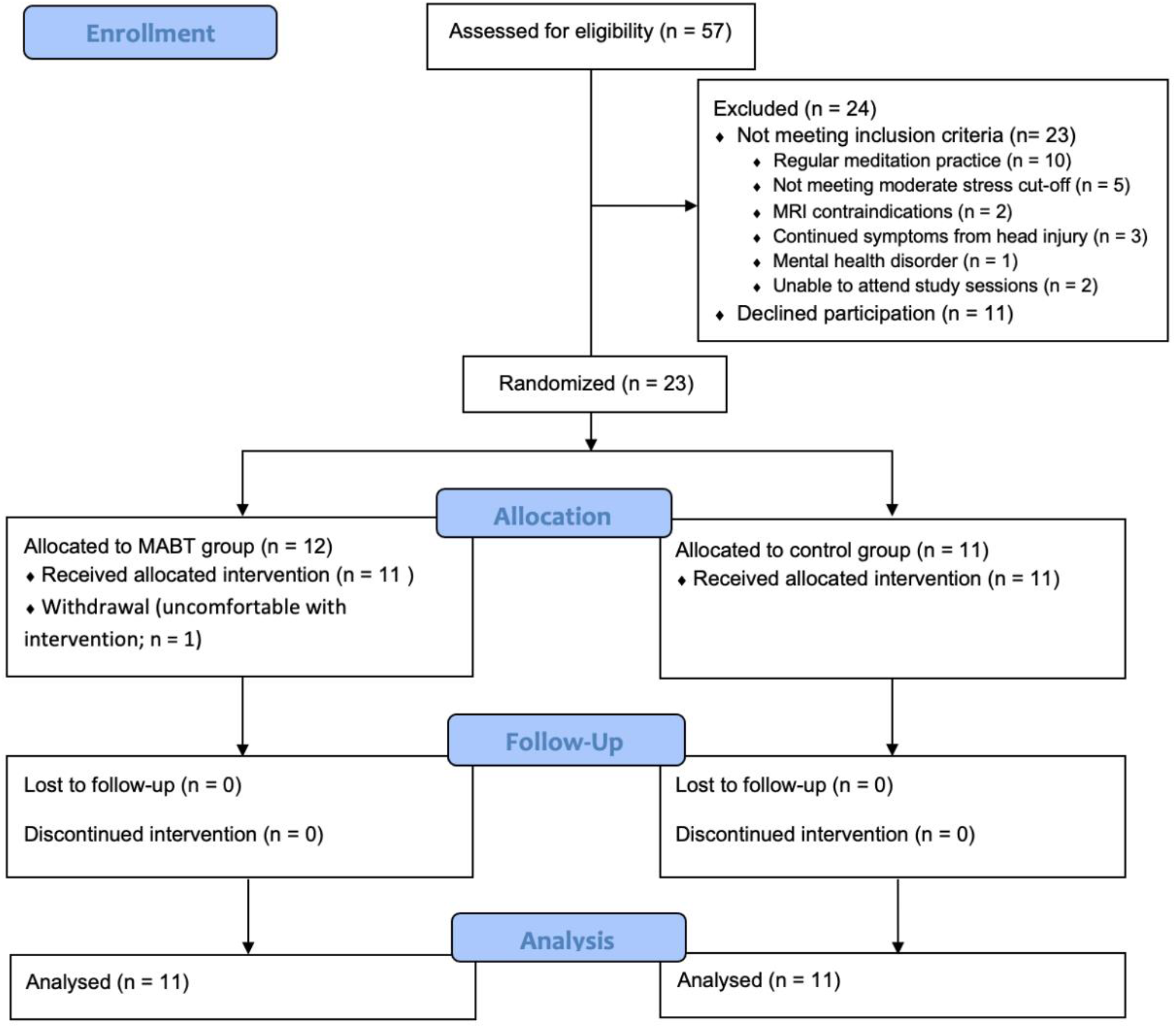
CONSORT flow diagram. Fifty-seven participants were recruited initially. Twenty-three participants were randomized to the MABT group (n = 12) and the control group (n = 11). Eleven participants from each group were included in data analysis.

**Supplementary Figure 2.**
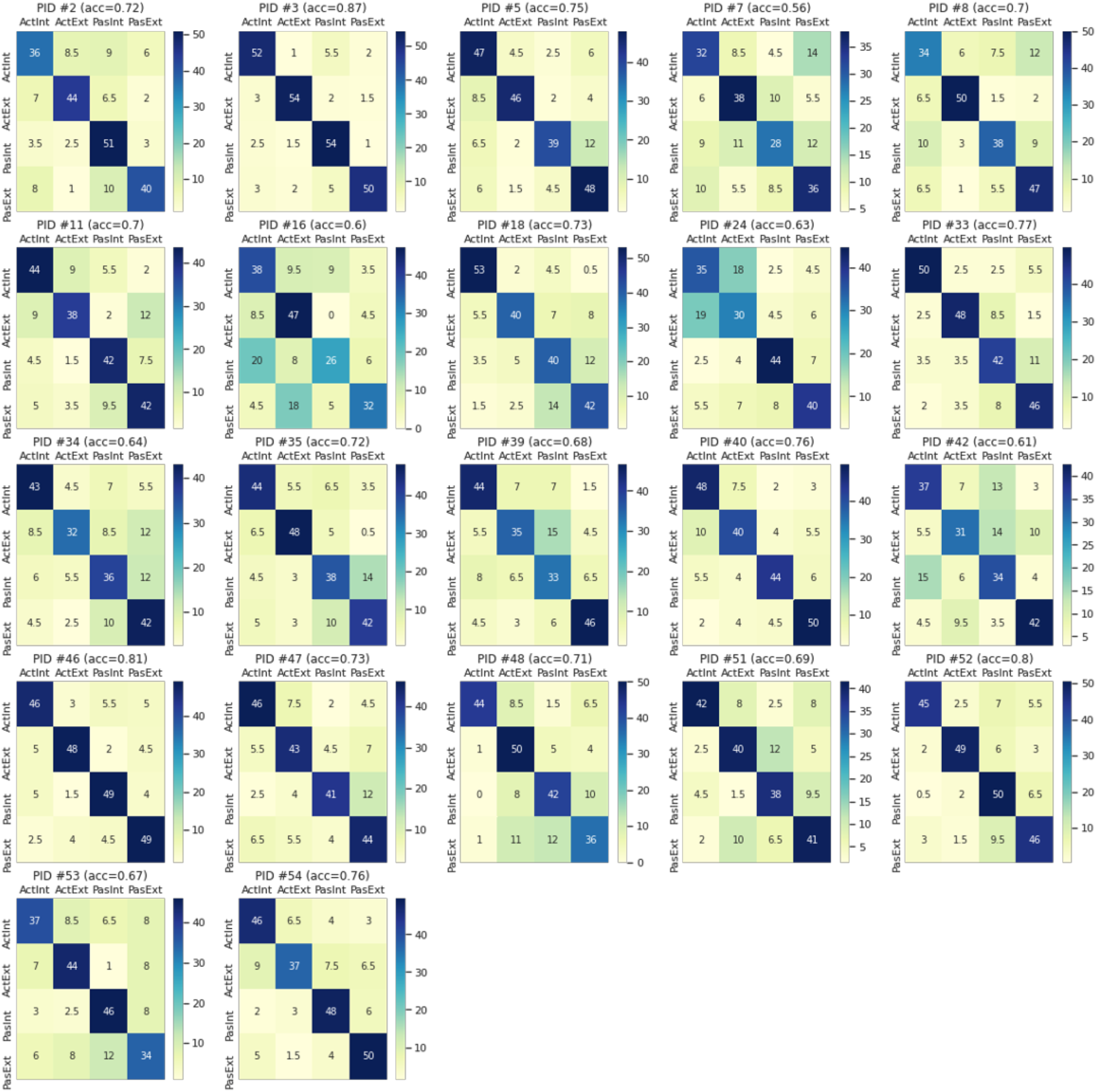
Individual participant within-session classification confusion matrices. These confusion matrices presented each participant’s classification accuracy across Active Interoception, Active Exteroception, Passive Interoception, and Passive Exteroception conditions. The dark diagonals in these matrices suggested that no mental state was systematically misclassified as another mental state.

**Supplementary Figure 3.**
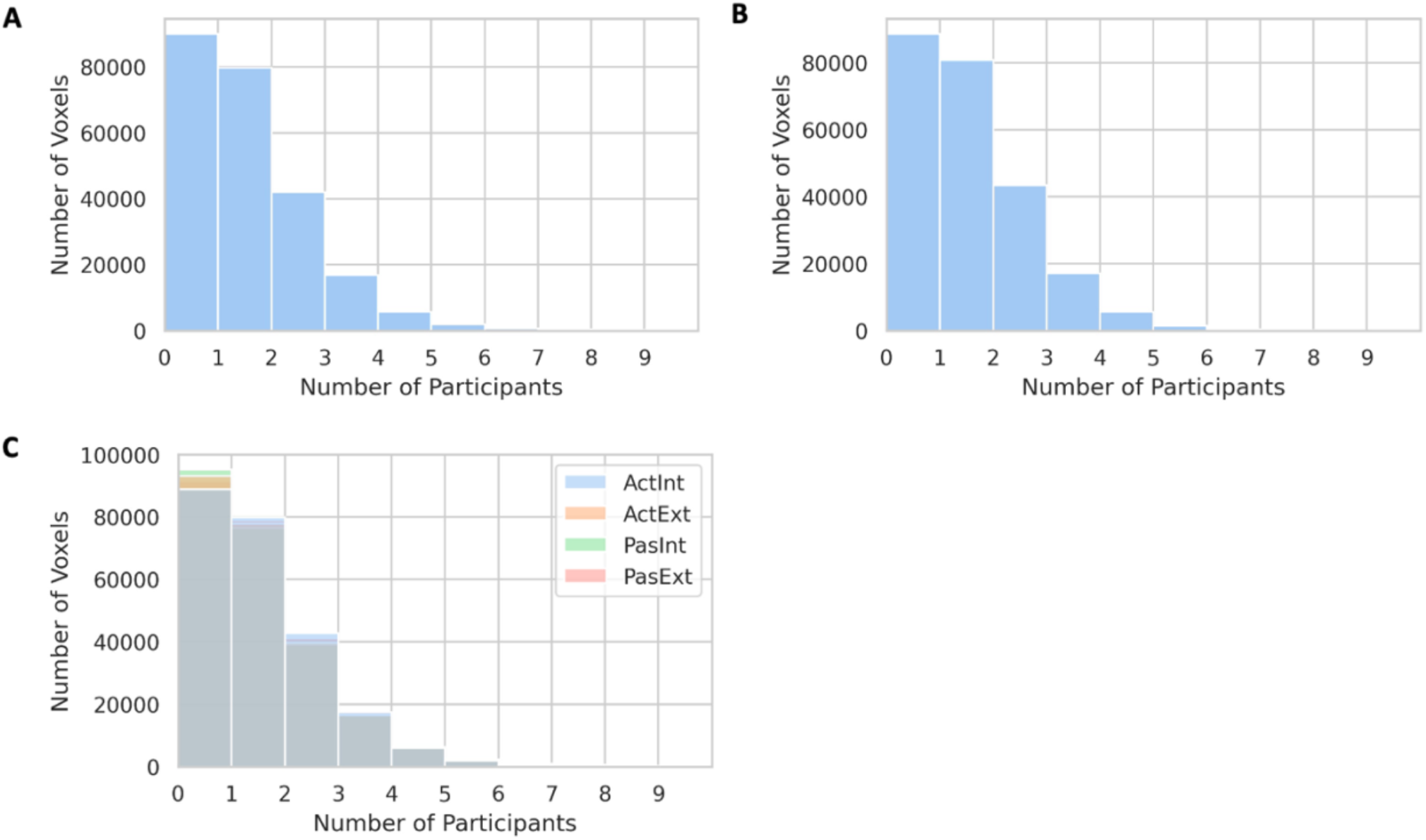
Shared important voxels in IEAT classification. Overall, very few voxels were important for more than 2 participants in the classification of attentional states. **(A)** Important voxels in the classification between Interoception and Exteroception. **(B)** Important voxels in the classification between Active Interoception and Active Exteroception. **(C)** Important voxels in the classification between Active Interoception, Active Exteroception, Passive Interoception, and Passive Exteroception.

**Supplementary Figure 4.**
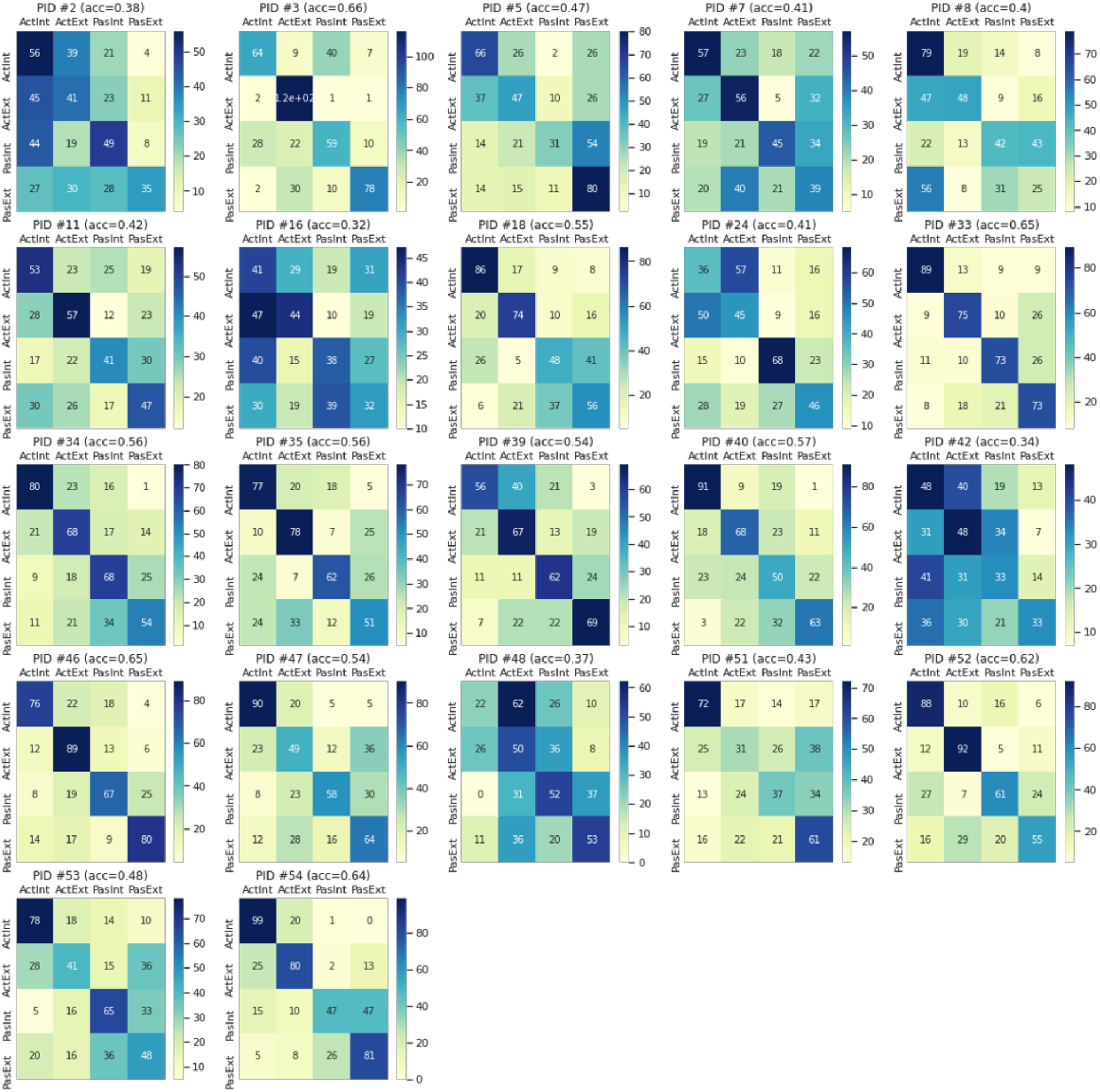
Individual participant across-session classification confusion matrices. These confusion matrices presented each participant’s classification accuracy across Active Interoception, Active Exteroception, Passive Interoception, and Passive Exteroception conditions. Compared to within-session classification, different mental states appeared more confusable in this analysis. For example, the mental states of participants #2 and #42 were classified more as active conditions.

**Supplementary Figure 5.**
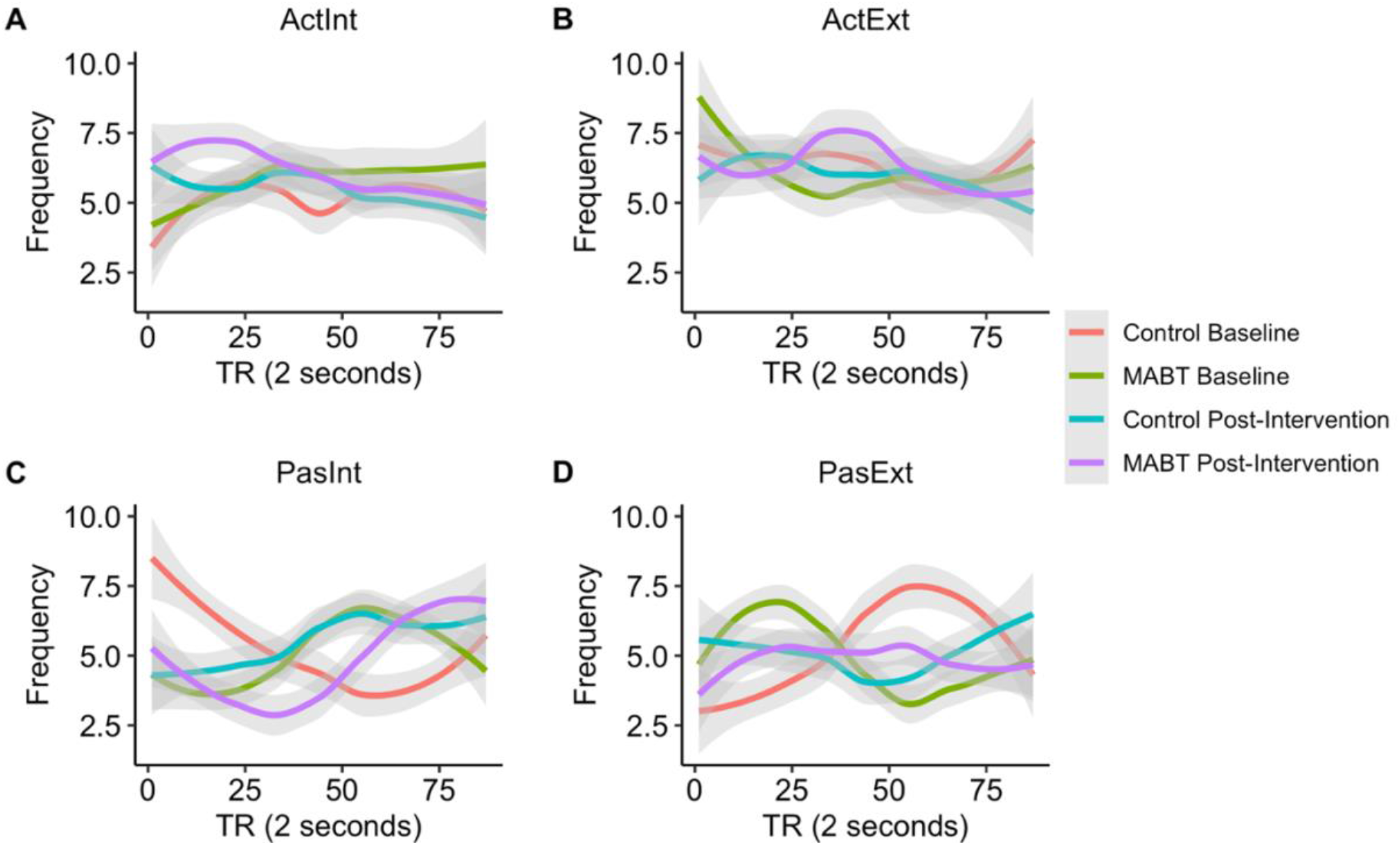
Timecourse of estimated frequency of mental states during the SIAT by Group and Session. For MABT and Control groups, sustained attention readouts were plotted separately for baseline and post-intervention sessions, for each of the four mental states: **(A)** Active Interoception, **(B)** Active Exteroception, **(C)** Passive Interoception, and **(D)** Passive Exteroception. Frequency here refers to the number of participants whose readout was in a certain mental state at each TR. **(A)** There was a significant Group × Session × TR interaction (β = −0.03, 95% CI [-0.06, 0], p = .049) for Active Interoception, likely driven by MABT post-intervention. **(C)** & **(D)** All interactions and main effects were significant in the two passive conditions.

**Supplementary Table 1.** Simple Effects of Active Interoception, Active Exteroception, Passive Interoception, and Passive Exteroception Readout Frequency Estimates.

| Active Interoception Frequency |  |  |  | Active Exteroception Frequency |  |  |
| --- | --- | --- | --- | --- | --- | --- |
| *Predictors* | *Estimates* | *CI* | *p* | *Estimates* | *CI* | *p* |
| (Intercept) | 4.94 | [4.15; 5.73] | **<0.001** | 6.69 | [5.83; 7.55] | **<0.001** |
| Group | -0.03 | [-1.15; 1.08] | 0.956 | 0.11 | [-1.10; 1.32] | 0.858 |
| Session | 1.03 | [-0.09; 2.14] | 0.071 | 0.21 | [-1.00; 1.42] | 0.733 |
| TR | 0.01 | [-0.01; 0.02] | 0.42 | -0.01 | [-0.03; 0.01] | 0.26 |
| Group * Session | 1.53 | [-0.05; 3.10] | 0.058 | -0.34 | [-2.05; 1.38] | 0.699 |
| Group * TR | 0.01 | [-0.01; 0.04] | 0.203 | -0.01 | [-0.03; 0.02] | 0.526 |
| Session * TR | -0.02 | [-0.04; 0.00] | 0.092 | -0.01 | [-0.03; 0.01] | 0.4 |
| Group * Session * TR | -0.03 | [-0.06; 0.00] | **0.049** | 0.02 | [-0.02; 0.05] | 0.352 |
| Observations | 348 |  |  | 348 |  |  |
| R<sup>2</sup> / R<sup>2</sup> adjusted | 0.099 / 0.080 |  |  | 0.038 / 0.018 |  |  |

| Passive Interoception Frequency |  |  |  | Passive Exteroception Frequency |  |  |
| --- | --- | --- | --- | --- | --- | --- |
| *Predictors* | *Estimates* | *CI* | *p* | *Estimates* | *CI* | *p* |
| (Intercept) | 6.94 | [6.13; 7.75] | **<0.001** | 3.43 | [2.60; 4.27] | **<0.001** |
| Group | -3.19 | [-4.34; -2.04] | **<0.001** | 3.11 | [1.93; 4.29] | **<0.001** |
| Session | -2.64 | [-3.79; -1.50] | **<0.001** | 1.41 | [0.23; 2.58] | **0.019** |
| TR | -0.04 | [-0.06; -0.03] | **<0.001** | 0.04 | [0.03; 0.06] | **<0.001** |
| Group * Session | 1.74 | [0.12; 3.36] | **0.036** | -2.93 | [-4.59; -1.26] | **0.001** |
| Group * TR | 0.07 | [0.05; 0.09] | **<0.001** | -0.08 | [-0.10; -0.05] | **<0.001** |
| Session * TR | 0.07 | [0.05; 0.09] | **<0.001** | -0.04 | [-0.06; -0.02] | **0.001** |
| Group * Session * TR | -0.05 | [-0.09; -0.02] | **0.001** | 0.07 | [0.04; 0.10] | **<0.001** |
| Observations | 348 |  |  | 348 |  |  |
| R<sup>2</sup> / R<sup>2</sup> adjusted | 0.200 / 0.183 |  |  | 0.124 / 0.106 |  |  |

*Note.* Bolded *p* values were less than .05.

**Supplementary Figure 6.**
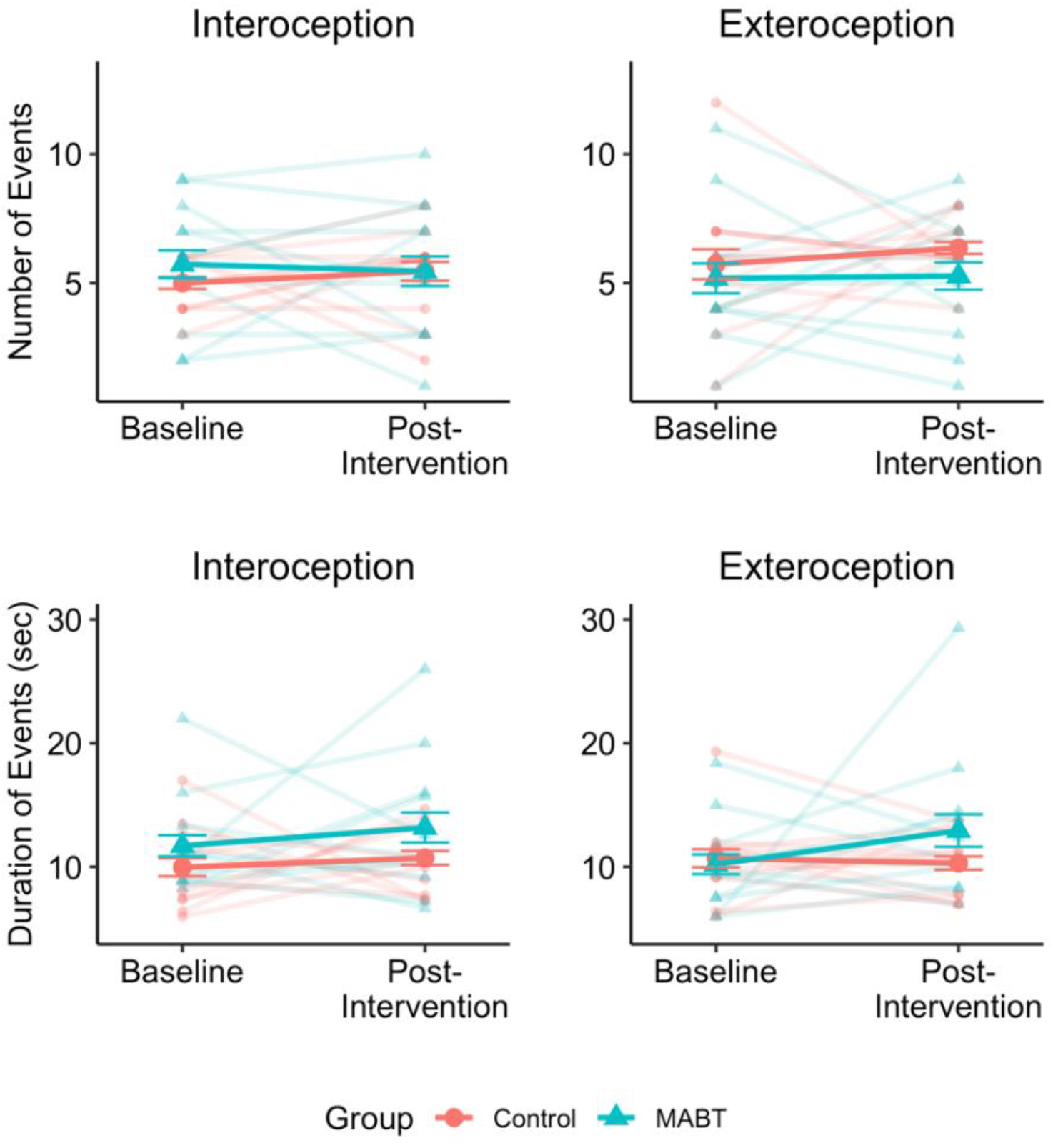
SIAT estimates by Group and Session. Using machine learning classifiers trained on the Active Interoception and Active Exteroception conditions, we predicted each participant’s moment-by-moment attention during the SIAT. The readouts were summarized into the number and average duration of interoceptive and exteroceptive events.

**Supplementary Table 2.**
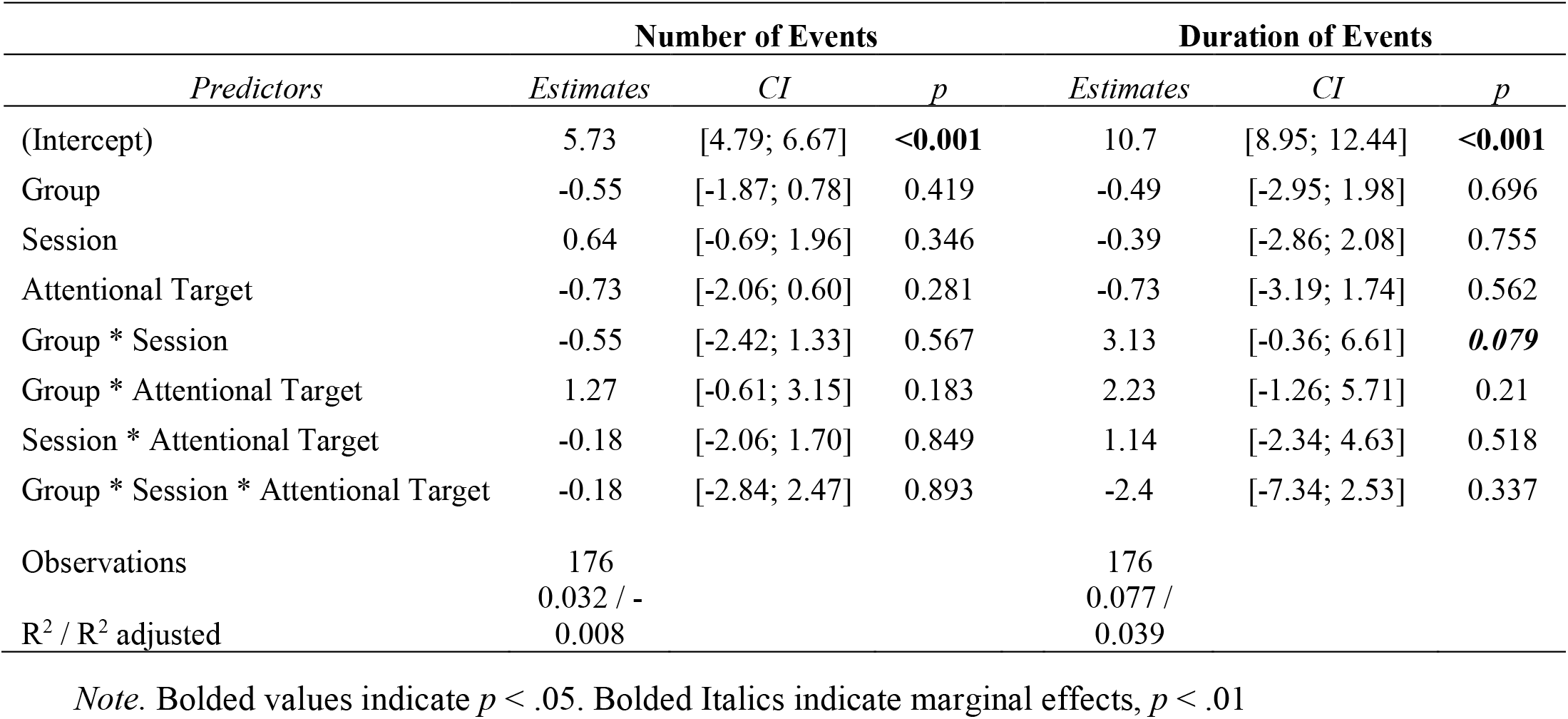
Modeling Attention Metrics Using Attentional Target, Group, and Session.

**Supplementary Figure 7.**
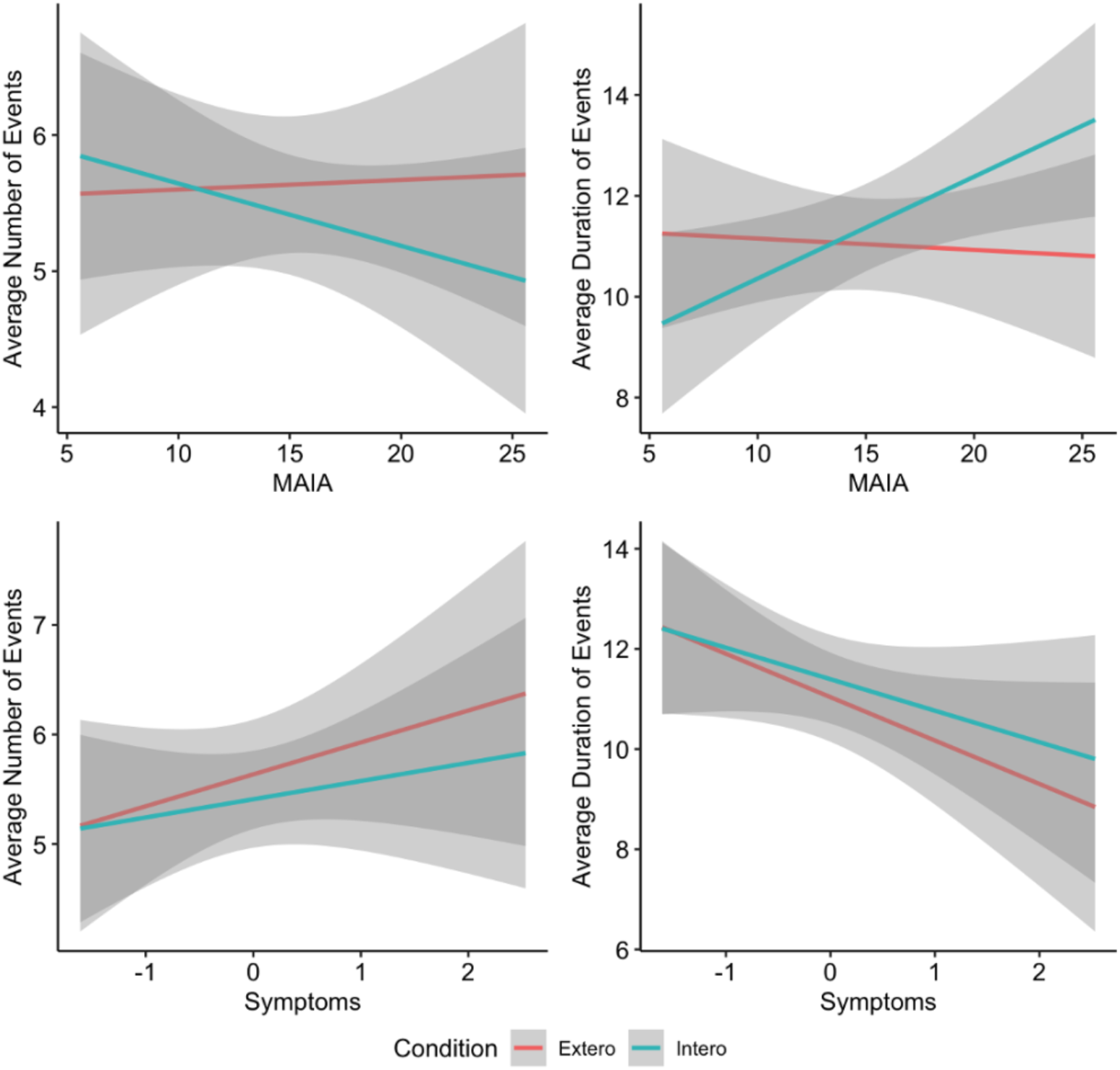
SIAT estimates and self-reported interoceptive awareness and affective symptom burden. MAIA and symptom scores were used to predict classifier model estimates of active interoceptive and exteroceptive attention during SIAT. The blue lines indicate interoceptive engagement metrics for the average number of events and the average duration of events. The red lines indicate exteroceptive engagement metrics.

**Supplementary Table 3.** Linear Mixed Model Output for MAIA/Symptom Ratings and Machine Learning Model Estimates (Trained on Active Interoception and Active Exteroception

| *Predictors* | **Number of Events** |  |  | **Average Duration of Events** |  |  |
| --- | --- | --- | --- | --- | --- | --- |
|  | *Estimates* | *CI* | *p* | *Estimates* | *CI* | *p* |
| (Intercept) | 5.29 | [3.27; 7.31] | **<0.001** | 11.36 | [7.56; 15.16] | **<0.001** |
| MAIA Rating | 0.02 | [-0.10; 0.15] | 0.716 | -0.02 | [-0.26; 0.22] | 0.861 |
| Attentional Target | 0.57 | [-2.06; 3.20] | 0.67 | -3.04 | [-8.38; 2.30] | 0.265 |
| MAIA Rating * Attentional Target | -0.05 | [-0.22; 0.11] | 0.528 | 0.22 | [-0.11; 0.56] | 0.187 |
| Random Effects |  |  |  |  |  |  |
| $\sigma^2$ | | 4.31 | | | 17.78 | |
| $\tau_{00}$ Subjects | | 0.81 | | | 0.19 | |
| ICC |  | 0.16 |  |  | 0.01 |  |
| N Subjects |  | 22 |  |  | 22 |  |
| Observations |  | 88 |  |  | 88 |  |
| Marginal R<sup>2</sup> / Conditional R<sup>2</sup> |  | 0.006 / 0.163 |  |  | 0.034 / 0.044 |  |
| *Predictors* | **Number of Events** |  |  | **Average Duration of Events** |  |  |
|  | *Estimates* | *CI* | *p* | *Estimates* | *CI* | *p* |
| (Intercept) | 5.64 | [4.92; 6.36] | **<0.001** | 11.04 | [9.78; 12.29] | **<0.001** |
| Symptom Rating | 0.31 | [-0.40; 1.02] | 0.388 | -0.87 | [-2.16; 0.42] | 0.188 |
| Attentional Target | -0.23 | [-1.09; 0.64] | 0.606 | 0.36 | [-1.41; 2.13] | 0.692 |
| Symptom Rating * Attentional Target | -0.13 | [-1.01; 0.76] | 0.783 | 0.24 | [-1.59; 2.06] | 0.798 |
| Random Effects |  |  |  |  |  |  |
| $\sigma^2$ | | 4.28 | | | 18 | |
| $\tau_{00}$ Subjects | | 0.83 | | | 0 | |
| ICC |  | 0.16 |  |  |  |  |
| N Subjects |  | 22 |  |  | 22 |  |
| Observations |  | 88 |  |  | 88 |  |
| Marginal R<sup>2</sup> / Conditional R<sup>2</sup> |  | 0.015 / 0.174 |  |  | 0.031 / NA |  |

*Note.* Attentional Target = exteroceptive vs. interoception attention. Bolded p values were less than .05.

**Supplementary Figure 8.**
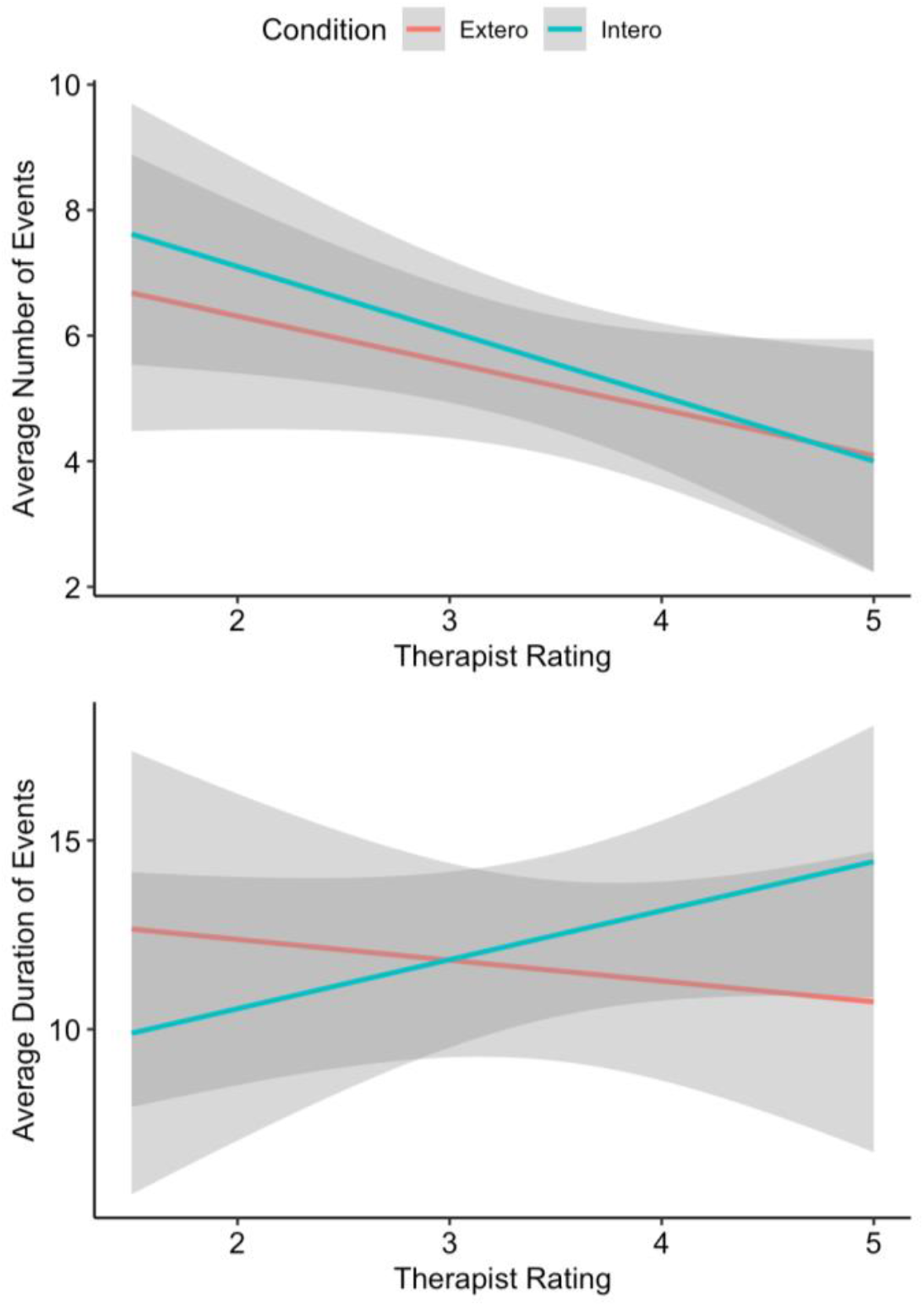
SIAT estimates and therapist-reported interoceptive awareness. For the MABT group only, therapist-rated interoceptive awareness (in MABT sessions 5 to 8) were used to predict classifier model estimates of active interoceptive and exteroceptive attention during SIAT. The blue lines indicate interoceptive engagement metrics for the average number of events and the average duration of events. The red lines indicate exteroceptive engagement metrics.

**Supplementary Table 4.** Linear Mixed Model Output for Therapist Rating and Machine Learning Model Estimates (Trained on Active Interoception and Active Exteroception Conditions).

| *Predictors* | **Number of Events** |  |  | **Average Duration of Events** |  |  |
| --- | --- | --- | --- | --- | --- | --- |
|  | *Estimates* | *CI* | *p* | *Estimates* | *CI* | *p* |
| (Intercept) | 7.79 | [4.26; 11.32] | **<0.001** | 13.48 | [6.40; 20.56] | **<0.001** |
| Therapist Rating | -0.74 | [-1.71; 0.23] | 0.129 | -0.55 | [-2.49; 1.39] | 0.569 |
| Condition | 1.37 | [-3.10; 5.85] | 0.538 | -5.53 | [-15.33; 4.26] | 0.260 |
| Therapist Rating * Condition | -0.29 | [-1.52; 0.93] | 0.633 | 1.85 | [-0.84; 4.53] | 0.171 |
| Random Effects |  |  |  |  |  |  |
| $\sigma^2$ | | | 5.36 | | | 25.69 |
| $\tau_{00}$ Subjects | | | 0.65 | | | 0.58 |
| ICC |  |  | 0.11 |  |  | 0.02 |
| N Subjects |  |  | 11 |  |  | 11 |
| Observations |  |  | 44 |  |  | 44 |
| Marginal R<sup>2</sup> / Conditional R<sup>2</sup> | 0.158 / 0.249 |  |  | 0.056 / 0.076 |  |  |

### Script for Audio-Guided Sustained Interoceptive Attention

“First, slide one of your hands so that the palm lies flat and firmly upon your upper chest. I’m going to lead you in a simple meditation involving focused attention on this area of your body. The scanner will turn on after I’m done talking. Then I’d like you to continue the meditation on your own for a couple of minutes. To begin, let your eyes close and let your thoughts quiet as you enter a state of meditation. I’d like to invite you to feel the weight of your hand firmly and gently pressing in and gently holding your upper chest. Allow your attention to come down into your chest underneath your hand as though you’re traveling down in your body into the space between your breastbone and your backbone. With your inner attention, notice the feeling of your body moving to the inhale and the exhale of your breath. Notice your sense of this place in your body and allow yourself to simply be here as though you’re keeping yourself company inside. Continue to let yourself attend to this area of your body for the next couple of minutes. If you start thinking or notice yourself drifting off, just bring your attention back to your hand and allow yourself to sink back down inside your chest. I’m going to stop talking now and you can continue this meditation and open your eyes only after you hear the scanner turn off.**”**

### Neuroimaging Pre-Processing Details

Preprocessing was performed using fMRIPrep 20.0.6 (Esteban, Markiewicz, et al. (2018); Esteban, Blair, et al. (2018); RRID:SCR_016216), which is based on Nipype 1.4.2 (Gorgolewski et al. (2011); Gorgolewski et al. (2018); RRID:SCR_002502).

#### Anatomical data preprocessing

A total of 2 T1-weighted (T1w) images were found within the input BIDS dataset. All of them were corrected for intensity non-uniformity (INU) with N4BiasFieldCorrection (Tustison et al. 2010), distributed with ANTs 2.2.0 (Avants et al. 2008, RRID:SCR_004757). The T1w-reference was then skull-stripped with a Nipype implementation of the antsBrainExtraction.sh workflow (from ANTs), using OASIS30ANTs as target template. Brain tissue segmentation of cerebrospinal fluid (CSF), white-matter (WM) and gray-matter (GM) was performed on the brain-extracted T1w using fast (FSL 5.0.9, RRID:SCR_002823, Zhang, Brady, and Smith 2001). A T1w-reference map was computed after registration of 2 T1w images (after INU-correction) using mri_robust_template (FreeSurfer 6.0.1, Reuter, Rosas, and Fischl 2010). Brain surfaces were reconstructed using recon-all (FreeSurfer 6.0.1, RRID:SCR_001847, Dale, Fischl, and Sereno 1999), and the brain mask estimated previously was refined with a custom variation of the method to reconcile ANTs-derived and FreeSurfer-derived segmentations of the cortical gray-matter of Mindboggle (RRID:SCR_002438, Klein et al. 2017). Volume-based spatial normalization to one standard space (MNI152NLin2009cAsym) was performed through nonlinear registration with antsRegistration (ANTs 2.2.0), using brain-extracted versions of both T1w reference and the T1w template. The following template was selected for spatial normalization: ICBM 152 Nonlinear Asymmetrical template version 2009c [Fonov et al. (2009), RRID:SCR_008796; TemplateFlow ID: MNI152NLin2009cAsym].

#### Functional data preprocessing

For each BOLD run found per subject (across all tasks and sessions), the following preprocessing was performed. First, a reference volume and its skull-stripped version were generated using a custom methodology of fMRIPrep. Susceptibility distortion correction (SDC) was omitted. The BOLD reference was then co-registered to the T1w reference using bbregister (FreeSurfer) which implements boundary-based registration (Greve and Fischl 2009). Co-registration was configured with six degrees of freedom. Head-motion parameters with respect to the BOLD reference (transformation matrices, and six corresponding rotation and translation parameters) are estimated before any spatiotemporal filtering using mcflirt (FSL 5.0.9, Jenkinson et al. 2002). The BOLD time-series (including slice-timing correction when applied) were resampled onto their original, native space by applying the transforms to correct for head-motion. These resampled BOLD time-series will be referred to as preprocessed BOLD in original space, or just preprocessed BOLD. The BOLD time-series were resampled into standard space, generating a preprocessed BOLD run in MNI152NLin2009cAsym space. First, a reference volume and its skull-stripped version were generated using a custom methodology of fMRIPrep. Several confounding time-series were calculated based on the preprocessed BOLD: framewise displacement (FD), DVARS and three region-wise global signals. FD and DVARS are calculated for each functional run, both using their implementations in Nipype (following the definitions by Power et al. 2014). The three global signals are extracted within the CSF, the WM, and the whole-brain masks. Additionally, a set of physiological regressors were extracted to allow for component-based noise correction (CompCor, Behzadi et al. 2007). Principal components are estimated after high-pass filtering the preprocessed BOLD time-series (using a discrete cosine filter with 128s cut-off) for the two CompCor variants: temporal (tCompCor) and anatomical (aCompCor). tCompCor components are then calculated from the top 5% variable voxels within a mask covering the subcortical regions. This subcortical mask is obtained by heavily eroding the brain mask, which ensures it does not include cortical GM regions. For aCompCor, components are calculated within the intersection of the aforementioned mask and the union of CSF and WM masks calculated in T1w space, after their projection to the native space of each functional run (using the inverse BOLD-to-T1w transformation). Components are also calculated separately within the WM and CSF masks. For each CompCor decomposition, the k components with the largest singular values are retained, such that the retained components’ time series are sufficient to explain 50 percent of variance across the nuisance mask (CSF, WM, combined, or temporal). The remaining components are dropped from consideration. The head-motion estimates calculated in the correction step were also placed within the corresponding confounds file. The confound time series derived from head motion estimates and global signals were expanded with the inclusion of temporal derivatives and quadratic terms for each (Satterthwaite et al. 2013). Frames that exceeded a threshold of 0.5 mm FD or 1.5 standardised DVARS were annotated as motion outliers. All resamplings can be performed with a single interpolation step by composing all the pertinent transformations (i.e. head-motion transform matrices, susceptibility distortion correction when available, and co-registrations to anatomical and output spaces). Gridded (volumetric) resamplings were performed using antsApplyTransforms (ANTs), configured with Lanczos interpolation to minimize the smoothing effects of other kernels (Lanczos 1964). Non-gridded (surface) resamplings were performed using mri_vol2surf (FreeSurfer).

Many internal operations of fMRIPrep use Nilearn 0.6.2 (Abraham et al. 2014, RRID:SCR_001362), mostly within the functional processing workflow. For more details of the pipeline, see the section corresponding to workflows in fMRIPrep’s documentation.

## References

Abraham, A., Pedregosa, F., Eickenberg, M., Gervais, P., Mueller, A., Kossaifi, J., Gramfort, A., Thirion, B., & Varoquaux, G. (2014). Machine learning for neuroimaging with scikit-learn. Frontiers in Neuroinformatics, 8. https://www.frontiersin.org/article/10.3389/fninf.2014.00014

Baltazar, M., Grezes, J., Geoffray, M., Picq, J., & Conty, L. (2021). Neural correlates of interoceptive accuracy: Beyond cardioception. European Journal of Neuroscience, 54(10), 7642–7653. https://doi.org/10.1111/ejn.15510

Barrett, L. F., & Quigley, K. S. (2021). Interoception: The Secret Ingredient. Cerebrum: The Dana Forum on Brain Science, 2021, cer-06-21.

Barsky, A. J., Peekna, H. M., & Borus, J. F. (2001). Somatic symptom reporting in women and men. Journal of General Internal Medicine, 16(4), 266–275. https://doi.org/10.1046/j.1525-1497.2001.00229.x

Bates, D., Mächler, M., Bolker, B., & Walker, S. (2015). Fitting linear mixed-effects models using lme4. Journal of Statistical Software, 67(1), 1–48. https://doi.org/10.18637/jss.v067.i01

Bosse, T., Jonker, C. M., & Treur, J. (2008). Formalization of Damasio’s theory of emotion, feeling and core consciousness. Consciousness and Cognition, 17(1), 94–113. https://doi.org/10.1016/j.concog.2007.06.006

Brown, K. W., & Cordon, S. (2009). Toward a Phenomenology of Mindfulness: Subjective Experience and Emotional Correlates. In F. Didonna (Ed.), Clinical Handbook of Mindfulness (pp. 59–81). Springer New York. https://doi.org/10.1007/978-0-387-09593-6_5

Chikazoe, J., Lee, D. H., Kriegeskorte, N., & Anderson, A. K. (2014). Population coding of affect across stimuli, modalities and individuals. Nature Neuroscience, 17(8), 1114–1122. https://doi.org/10.1038/nn.3749

Clark, I. A., Niehaus, K. E., Duff, E. P., Di Simplicio, M. C., Clifford, G. D., Smith, S. M., Mackay, C. E., Woolrich, M. W., & Holmes, E. A. (2014). First steps in using machine learning on fMRI data to predict intrusive memories of traumatic film footage. Behaviour Research and Therapy, 62, 37–46. https://doi.org/10.1016/j.brat.2014.07.010

Cohen, S., Kamarck, T., & Mermelstein, R. (1983). A global measure of perceived stress. Journal of Health and Social Behavior, 24(4), 385. https://doi.org/10.2307/2136404

Craig, A. D. (2002). How do you feel? Interoception: The sense of the physiological condition of the body. Nature Reviews Neuroscience, 3(8), 655–666. https://doi.org/10.1038/nrn894

Craig, A. D. (2003). Interoception: The sense of the physiological condition of the body. Curr Opin Neurobiol, 13(4), 500–505. https://doi.org/S0959438803000904 [pii]

Critchley, H. D., & Harrison, N. A. (2013). Visceral influences on brain and behavior. Neuron, 77(4), 624–638. https://doi.org/10.1016/j.neuron.2013.02.008

Davatzikos, C. (2019). Machine learning in neuroimaging: Progress and challenges. NeuroImage, 197, 652–656. https://doi.org/10.1016/j.neuroimage.2018.10.003

de Jong, M., Lazar, S. W., Hug, K., Mehling, W. E., Hölzel, B. K., Sack, A. T., Peeters, F., Ashih, H., Mischoulon, D., & Gard, T. (2016). Effects of mindfulness-based cognitive therapy on body awareness in patients with chronic pain and comorbid depression. Frontiers in Psychology, 7. https://doi.org/10.3389/fpsyg.2016.00967

Dixon, M. L., De La Vega, A., Mills, C., Andrews-Hanna, J., Spreng, R. N., Cole, M. W., & Christoff, K. (2018). Heterogeneity within the frontoparietal control network and its relationship to the default and dorsal attention networks. Proceedings of the National Academy of Sciences, 115(7). https://doi.org/10.1073/pnas.1715766115

Domschke, K., Stevens, S., Pfleiderer, B., & Gerlach, A. L. (2010). Interoceptive sensitivity in anxiety and anxiety disorders: An overview and integration of neurobiological findings. Clinical Psychology Review, 30(1), 1–11.

Dunn, B. D., Dalgleish, T., Ogilvie, A. D., & Lawrence, A. D. (2007). Heartbeat perception in depression. Behaviour Research and Therapy, 45(8), 1921–1930. https://doi.org/10.1016/j.brat.2006.09.008

Dunn, B. D., Galton, H. C., Morgan, R., Evans, D., Oliver, C., Meyer, M., Cusack, R., Lawrence, A. D., & Dalgleish, T. (2010). Listening to your heart. How interoception shapes emotion experience and intuitive decision making. Psychol Sci, 21(12), 1835–1844. https://doi.org/10.1177/0956797610389191

Ellsworth, P. C. (2013). Appraisal Theory: Old and New Questions. Emotion Review, 5(2), 125–131. https://doi.org/10.1177/1754073912463617

Farb, N. A. S., Anderson, A. K., Mayberg, H., Bean, J., McKeon, D., & Segal, Z. V. (2010). Minding one’s emotions: Mindfulness training alters the neural expression of sadness. Emotion, 10(1), 25–33. https://doi.org/10.1037/a0017151

Farb, N. A. S., Daubenmier, J., Price, C. J., Gard, T., Kerr, C., Dunn, B. D., Klein, A. C., Paulus, M. P., & Mehling, W. E. (2015). Interoception, contemplative practice, and health. Frontiers in Psychology, 6. https://doi.org/10.3389/fpsyg.2015.00763

Farb, N. A. S., Segal, Z. V., Mayberg, H., Bean, J., McKeon, D., Fatima, Z., & Anderson, A. K. (2007). Attending to the present: Mindfulness meditation reveals distinct neural modes of self-reference. Social Cognitive and Affective Neuroscience, 2(4), 313–322. https://doi.org/10.1093/scan/nsm030

Farb, N. A. S., Zuo, Z., & Price, C. (2022). *Neural Dynamics of Interoceptive Attention and Awareness: A Within-Participant fMRI Study* (p. 2022.05.27.493743). bioRxiv. https://www.biorxiv.org/content/10.1101/2022.05.27.493743v1

Farb, N. A., Segal, Z. V., & Anderson, A. K. (2013). Attentional modulation of primary interoceptive and exteroceptive cortices. Cereb Cortex, 23(1), 114–126. https://doi.org/10.1093/cercor/bhr385

Flynn, F. G. (1999). Anatomy of the insula functional and clinical correlates. Aphasiology, 13(1), 55–78. https://doi.org/10.1080/026870399402325

Furman, D. J., Waugh, C. E., Bhattacharjee, K., Thompson, R. J., & Gotlib, I. H. (2013). Interoceptive awareness, positive affect, and decision making in Major Depressive Disorder. Journal of Affective Disorders, 151(2), 780–785. https://doi.org/10.1016/j.jad.2013.06.044

Glenn, D. E., Acheson, D. T., Geyer, M. A., Nievergelt, C. M., Baker, D. G., & Risbrough, V. B. (2016). High and low threshold for startle reactivity associated with PTSD symptoms but not PTSD risk: Evidence from a prospective study of active duty marines. Depression and Anxiety, 33(3), 192–202. https://doi.org/10.1002/da.22475

Greenlee, M. W., & Tse, P. U. (2008). Functional neuroanatomy of the human visual system: A review of functional MRI studies. In B. Lorenz & F.-X. Borruat (Eds.), Pediatric Ophthalmology, Neuro-Ophthalmology, Genetics (pp. 119–138). Springer. https://doi.org/10.1007/978-3-540-33679-2_8

Haxby, J. V. (2012). Multivariate pattern analysis of fMRI: The early beginnings. NeuroImage, 62(2), 852–855. https://doi.org/10.1016/j.neuroimage.2012.03.016

Herbert, B. M., & Pollatos, O. (2014). Attenuated interoceptive sensitivity in overweight and obese individuals. Eating Behaviors, 15(3), 445–448.

Jabakhanji, R., Vigotsky, A. D., Bielefeld, J., Huang, L., Baliki, M. N., Iannetti, G. D., & Apkarian, A. V. (2022). Limits of decoding mental states with fMRI. Cortex. https://doi.org/10.1016/j.cortex.2021.12.015

Khalsa, S. S., Adolphs, R., Cameron, O. G., Critchley, H. D., Davenport, P. W., Feinstein, J. S., Feusner, J. D., Garfinkel, S. N., Lane, R. D., Mehling, W. E., Meuret, A. E., Nemeroff, C. B., Oppenheimer, S., Petzschner, F. H., Pollatos, O., Rhudy, J. L., Schramm, L. P., Simmons, W. K., Stein, M. B.,. Interoception Summit 2016 participants. (2018). Interoception and Mental Health: A Roadmap. Biological Psychiatry. Cognitive Neuroscience and Neuroimaging, 3(6), 501–513. https://doi.org/10.1016/j.bpsc.2017.12.004

Khalsa, S. S., Rudrauf, D., Feinstein, J. S., & Tranel, D. (2009). The pathways of interoceptive awareness. Nat Neurosci, 12(12), 1494–1496. https://doi.org/10.1038/nn.2411

Kroenke, K., Spitzer, R. L., Williams, J. B. W., & Löwe, B. (2010). The Patient Health Questionnaire Somatic, Anxiety, and Depressive Symptom Scales: A systematic review. General Hospital Psychiatry, 32(4), 345–359. https://doi.org/10.1016/j.genhosppsych.2010.03.006

Kumar, M., Anderson, M. J., Antony, J. W., Baldassano, C., Brooks, P. P., Cai, M. B., Chen, P.-H. C., Ellis, C. T., Henselman-Petrusek, G., Huberdeau, D., Hutchinson, J. B., Li, Y. P., Lu, Q., Manning, J. R., Mennen, A. C., Nastase, S. A., Richard, H., Schapiro, A. C., Schuck, N. W.,. Norman, K. A. (2022). BrainIAK: The Brain Imaging Analysis Kit. Aperture Neuro, 2021(4), 42. https://doi.org/10.52294/31bb5b68-2184-411b-8c00-a1dacb61e1da

Kumar, M., Ellis, C. T., Lu, Q., Zhang, H., Capota, M., Willke, T. L., Ramadge, P. J., Turk-Browne, N. B., & Norman, K. A. (2020). BrainIAK tutorials: User-friendly learning materials for advanced fMRI analysis. PLOS Computational Biology, 16(1), e1007549. https://doi.org/10.1371/journal.pcbi.1007549

Lee, E.-H. (2012). Review of the psychometric evidence of the perceived stress scale. Asian Nursing Research, 6(4), 121–127. https://doi.org/10.1016/j.anr.2012.08.004

Li, D., Zucker, N. L., Kragel, P. A., Covington, V. E., & LaBar, K. S. (2017). Adolescent development of insula-dependent interoceptive regulation. Developmental Science, 20(5), e12438. https://doi.org/10.1111/desc.12438

Limanowski, J., & Blankenburg, F. (2013). Minimal self-models and the free energy principle. Frontiers in Human Neuroscience, 7.

Lotze, M., Erb, M., Flor, H., Huelsmann, E., Godde, B., & Grodd, W. (2000). FMRI evaluation of somatotopic representation in human primary motor cortex. NeuroImage, 11(5 Pt 1), 473-481. https://doi.org/10.1006/nimg.2000.0556

Matsumoto, R., Kitabayashi, Y., Narumoto, J., Wada, Y., Okamoto, A., Ushijima, Y., Yokoyama, C., Yamashita, T., Takahashi, H., Yasuno, F., Suhara, T., & Fukui, K. (2006). Regional cerebral blood flow changes associated with interoceptive awareness in the recovery process of anorexia nervosa. Progress in Neuro-Psychopharmacology & Biological Psychiatry, 30(7), 1265–1270. https://doi.org/10.1016/j.pnpbp.2006.03.042

Medford, N., & Critchley, H. D. (2010). Conjoint activity of anterior insular and anterior cingulate cortex: Awareness and response. Brain Structure and Function, 214(5), 535–549. https://doi.org/10.1007/s00429-010-0265-x

Mehling, W. E., Price, C., Daubenmier, J. J., Acree, M., Bartmess, E., & Stewart, A. (2012). The Multidimensional Assessment of Interoceptive Awareness (MAIA). PLoS ONE, 7(11), e48230. https://doi.org/10.1371/journal.pone.0048230

Mitchell, T. M., Hutchinson, R., Just, M. A., Niculescu, R. S., Pereira, F., & Wang, X. (2003). Classifying Instantaneous Cognitive States from fMRI Data. AMIA Annual Symposium Proceedings, 2003, 465–469.

Naqvi, N. H., & Bechara, A. (2010). The insula and drug addiction: An interoceptive view of pleasure, urges, and decision-making. Brain Structure & Function, 214(0), 435–450. https://doi.org/10.1007/s00429-010-0268-7

Navarro-Haro, M. V., Modrego-Alarcón, M., Hoffman, H. G., López-Montoyo, A., Navarro-Gil, M., Montero-Marin, J., García-Palacios, A., Borao, L., & García-Campayo, J. (2019). Evaluation of a mindfulness-based intervention with and without Virtual Reality Dialectical Behavior Therapy® mindfulness Skills Training for the Treatment of Generalized Anxiety Disorder in Primary Care: A Pilot Study. Frontiers in Psychology, 10, 55. https://doi.org/10.3389/fpsyg.2019.00055

Norman, K. A., Polyn, S. M., Detre, G. J., & Haxby, J. V. (2006). Beyond mind-reading: Multi-voxel pattern analysis of fMRI data. Trends in Cognitive Sciences, 10(9), 424–430. https://doi.org/10.1016/j.tics.2006.07.005

Pedregosa, F., Varoquaux, G., Gramfort, A., Michel, V., Thirion, B., Grisel, O., Blondel, M., Prettenhofer, P., Weiss, R., Dubourg, V., Vanderplas, J., Passos, A., Cournapeau, D., Brucher, M., Perrot, M., & Duchesnay, É. (2011). Scikit-learn: Machine learning in Python. Journal of Machine Learning Research, 12(85), 2825–2830.

Pollatos, O., Füstös, J., & Critchley, H. D. (2012). On the generalised embodiment of pain: How interoceptive sensitivity modulates cutaneous pain perception. Pain, 153(8), 1680–1686.

Price, C. J., & Hooven, C. (2018). Interoceptive awareness skills for emotion regulation: Theory and approach of Mindful Awareness in Body-Oriented Therapy (MABT). Frontiers in Psychology, 9. https://doi.org/10.3389/fpsyg.2018.00798

Price, C. J., Merrill, J. O., McCarty, R. L., Pike, K. C., & Tsui, J. I. (2020). A pilot study of mindful body awareness training as an adjunct to office-based medication treatment of opioid use disorder. Journal of Substance Abuse Treatment, 108, 123–128. https://doi.org/10.1016/j.jsat.2019.05.013

Price, C. J., Thompson, E. A., Crowell, S. E., Pike, K., Cheng, S. C., Parent, S., & Hooven, C. (2019). Immediate effects of interoceptive awareness training through Mindful Awareness in Body-oriented Therapy (MABT) for women in substance use disorder treatment. Substance Abuse, 40(1), 102–115. https://doi.org/10.1080/08897077.2018.1488335

Price, C. J., Wells, E. A., Donovan, D. M., & Rue, T. (2012). Mindful awareness in body-oriented therapy as an adjunct to women’s substance use disorder treatment: A pilot feasibility study. Journal of Substance Abuse Treatment, 43(1), 94–107. https://doi.org/10.1016/j.jsat.2011.09.016

Quadt, L., Critchley, H. D., & Garfinkel, S. N. (2018). The neurobiology of interoception in health and disease: Neuroscience of interoception. Annals of the New York Academy of Sciences, 1428(1), 112–128. https://doi.org/10.1111/nyas.13915

Quigley, K. S., Kanoski, S., Grill, W. M., Barrett, L. F., & Tsakiris, M. (2021). Functions of Interoception: From Energy Regulation to Experience of the Self. Trends in Neurosciences, 44(1), 29–38. https://doi.org/10.1016/j.tins.2020.09.008

Raichle, M. E., & Snyder, A. Z. (2007). A default mode of brain function: A brief history of an evolving idea. Neuroimage, 37(4), 1083–1090; discussion 1097-9. https://doi.org/10.1016/j.neuroimage.2007.02.041

Rao, S. M., Binder, J. R., Hammeke, T. A., Bandettini, P. A., Bobholz, J. A., Frost, J. A., Myklebust, B. M., Jacobson, R. D., & Hyde, J. S. (1995). Somatotopic mapping of the human primary motor cortex with functional magnetic resonance imaging. Neurology, 45(5), 919–924. https://doi.org/10.1212/WNL.45.5.919

Seeley, W. W. (2019). The Salience Network: A Neural System for Perceiving and Responding to Homeostatic Demands. The Journal of Neuroscience: The Official Journal of the Society for Neuroscience, 39(50), 9878–9882. https://doi.org/10.1523/JNEUROSCI.1138-17.2019

Seeley, W. W., Menon, V., Schatzberg, A. F., Keller, J., Glover, G. H., Kenna, H., Reiss, A. L., & Greicius, M. D. (2007). Dissociable Intrinsic Connectivity Networks for Salience Processing and Executive Control. Journal of Neuroscience, 27(9), 2349–2356. https://doi.org/10.1523/JNEUROSCI.5587-06.2007

Seth, A. K., Suzuki, K., & Critchley, H. D. (2012). An interoceptive predictive coding model of conscious presence. Frontiers in Psychology, 2. https://doi.org/10.3389/fpsyg.2011.00395

Siemer, M., & Reisenzein, R. (2007). The process of emotion inference. Emotion, 7(1), 1–20. https://doi.org/10.1037/1528-3542.7.1.1

Smith, S. M. (2022). The affectively embodied perspective of the subject. Philosophical Psychology, 1-30. https://doi.org/10.1080/09515089.2022.2081143

Strigo, I. A., & Craig, A. D. B. (2016). Interoception, homeostatic emotions and sympathovagal balance. *Philosophical Transactions of the Royal Society of London. Series B*, Biological Sciences, 371(1708), 20160010. https://doi.org/10.1098/rstb.2016.0010

Szczepanski, S. M., Pinsk, M. A., Douglas, M. M., Kastner, S., & Saalmann, Y. B. (2013). Functional and structural architecture of the human dorsal frontoparietal attention network. Proceedings of the National Academy of Sciences, 110(39), 15806–15811. https://doi.org/10.1073/pnas.1313903110

Terhaar, J., Viola, F. C., Bär, K.-J., & Debener, S. (2012). Heartbeat evoked potentials mirror altered body perception in depressed patients. Clinical Neurophysiology, 123(10), 1950–1957. https://doi.org/10.1016/j.clinph.2012.02.086

Tsakiris, M., & Critchley, H. D. (2016). Interoception beyond homeostasis: Affect, cognition and mental health. Philosophical Transactions of the Royal Society B: Biological Sciences, 371(1708), 20160002. https://doi.org/10.1098/rstb.2016.0002

Tsakiris, M., Tajadura-Jimenez, A., & Costantini, M. (2011). Just a heartbeat away from one’s body: Interoceptive sensitivity predicts malleability of body-representations. Proceedings of the Royal Society B: Biological Sciences, 278(1717), 2470–2476.

Waskom, M. L. (2021). seaborn: Statistical data visualization. Journal of Open Source Software, 6(60), 3021. https://doi.org/10.21105/joss.03021

Weng, H. Y., Lewis-Peacock, J. A., Hecht, F. M., Uncapher, M. R., Ziegler, D. A., Farb, N. A. S., Goldman, V., Skinner, S., Duncan, L. G., Chao, M. T., & Gazzaley, A. (2020). Focus on the breath: Brain decoding reveals internal states of attention during meditation. Frontiers in Human Neuroscience, 14. https://doi.org/10.3389/fnhum.2020.00336

Wickham, H. (2016). *ggplot2: Elegant Graphics for Data Analysis* (2nd ed.). Springer International Publishing. https://doi.org/10.1007/978-3-319-24277-4

Wiens, S. (2005). Interoception in emotional experience. Curr Opin Neurol, 18(4), 442–447.

